# Probiotics supplementation to adult human small intestinal stoma microbiota causes dynamic increase in the community resistance to perturbations and nutrient utilization

**DOI:** 10.1101/2023.01.29.525861

**Authors:** Jack Jansma, Nicola U. Thome, Markus Schwalbe, Anastasia Chrysovalantou Chatziioannou, Somayah S. Elsayed, Gilles P. van Wezel, Pieter van den Abbeele, Saskia van Hemert, Sahar El Aidy

**Author notes:** Present address: Department of Chemistry, Laboratory of Analytical Biochemistry, University of Crete, Heraklion, Greece (Anastasia Chrysovalantou Chatziioannou). Corresponding author: Sahar El Aidy. Host-microbe Interactions, Groningen Biomolecular Sciences and Biotechnology Institute (GBB), University of Groningen, Nijenborgh 7, 9747 AG Groningen, The Netherlands., E.

## Abstract

The gut microbiota plays a pivotal role in health and disease. The use of probiotics as microbiota-targeted therapies is a promising strategy to improve host health. However, dynamic molecular mechanisms are often not elucidated, especially when targeting the small intestinal microbiota. Here, we show that supplementation of a probiotic formula (Ecologic®825) to the adult human small intestinal ileostoma microbiota counteracts the growth of *Enterococcaceae* and *Enterobacteriaceae* and reduces ethanol production, leading to major changes in nutrient utilization and resistance to perturbations. The observed alterations coincided with an initial increase in lactate production and decrease in pH by the probiotics, followed by a sharp increase in the levels of butyrate and propionate. Additionally, increased production of multiple *N*-acyl amino acids was detected in the stoma samples supplemented with the probiotic formula. Overall, this study shows how network theory can be used to improve the current or identify novel microbiota-targeted therapies. The outcome may help further understand the reported effects of these probiotic formula on the host.

## Introduction

The gastrointestinal tract of healthy humans harbors a distinct microbial community containing hundreds of bacterial species [1]. The stability, resilience, and resistance to perturbations of a microbial community is dependent on the microbiota composition and the interactions among each other as well as with the host [2, 3]. Often, these interactions are mediated by microbiota-produced metabolites resulting in metabolic interaction networks [4, 5]. Subsequently, disruption of these networks by medication, disease or dietary interventions may alter the function of the community with consequences on the host health [6]. For example, Sung et al. combined metagenomics sequencing data with experimentally validated metabolic transport and macromolecular degradation reactions, to construct and compare a metabolic interaction network of healthy controls and type 2 diabetic patients. Using this network approach, the authors showed alterations in the produced metabolites as well as in the species with the most significant influence on the network [7]. Other metabolic network approaches have been used to investigate the influence of the microbiota in diseases such as inflammatory bowel disease, obesity and Parkinson’s disease [8, 9]. Accordingly, network approaches can be used as a knowledge driven approach to identify novel microbiota-targeted therapies [10]. For example, Steinway et al. used a dynamic microbial interaction network approach to identify *Barnesiella intestinehominis* as a promising candidate for the treatment of *Clostridium difficile* infection [11].

Probiotics are defined as live microorganisms that, when administered in adequate amounts, confer a health benefit on the host [12]. Currently, probiotics are used in prevention and treatment of several disorders [13, 14]. It has been widely accepted that the impact of multi strain probiotic formulas instead of single strain probiotics is potentially greater on the host since the strains within the formula can complement each other [15]. Indeed, oral administration of a probiotic formula (Ecologic® 825) to healthy volunteers resulted in a minor shift in the fecal microbiota [16], while when administered orally to patients with irritable bowel syndrome, an increase in fecal levels of acetate and butyrate, accompanied by a reduction in fecal zonulin and symptom severity were observed [17]. Administration of this probiotic formula together with a prebiotic (Ecologic® 825/FOS P6) to healthy volunteers increased stool frequency, but did not affect zonulin levels and gastrointestinal symptoms [18]. The reported effects of the supplemented probiotics could be a result of their interaction with the resident microbiota. Thus, a better understanding of the molecular mechanisms that regulate the metabolic interactions between the different strains within a probiotic formula as well as between the probiotic formula and the residing microbiota is needed.

Another caveat in the probiotic research is the limited investigation of the small intestine, despite being a pivotal target for the probiotic activity [19]. Sampling the small intestine in humans is challenging, which hampers our understanding of the microbiota, diet and host interactions as well as small intestinal disorders [20]. Besides, the small intestinal environment is highly dynamic, with oxygen, pH and nutrient gradients, making the investigation of probiotic therapies targeting the small intestine difficult [21]. However, the use of stoma effluent from ileostomy subjects has been shown to serve as a surrogate for the *in vivo* small intestine [22–24].

In this study, we employed a dynamic correlation-based metabolic network approach combined with a multivariate analysis approach using experimental data obtained from proton-nuclear magnetic resonance (^1^H-NMR), liquid chromatography-mass spectrometry (LC-MS), shallow shotgun sequencing and flow cytometry performed on ileostoma samples grown using ex vivo SIFR® technology [25] to investigate the alterations caused by the supplementation of a 9-species probiotic formula.

## Material and methods

### Product supplementation to the ileostoma effluent

Ecologic® 825 (Winclove probiotics, The Netherlands), a probiotic formula, consisting of 9 different strains; *Bifidobacterium bifidum* W23, *Bifidobacterium lactis* W51, *Bifidobacterium lactis* W52, *Lactobacillus acidophilus* W22, *Lactobacillus casei* W56, *Lactobacillus paracasei* W20, *Lactobacillus plantarum* W62 *Lactobacillus salivarius* W24 and *Lactococcus lactis* W19, was tested at a dose of 10 CFU/mL. Prior to the introduction of the formula in the bioreactors, the microbial cells were washed by centrifugation and resuspension in anaerobic PBS.

### Fermentation experiment using SIFR® technology

Individual bioreactors were processed in parallel in a bioreactor management device (Cryptobiotix, Ghent, Belgium). Each bioreactor contained 5 mL of nutritional medium (M0014, Cryptobiotix, Ghent, Belgium) - microbial inoculum blend with or without supplementation of the probiotic formula (at 10^7^ CFU probiotic/mL), then sealed individually, before being rendered anaerobic. At the start of the incubation, oxygen was introduced at pO_2_ = 15 mmHg, thus providing relevant intraluminal oxygen levels for the proximal ileum [26]. After preparation, bioreactors were incubated under continuous agitation (140 rpm) at 37 °C for 24h (MaxQ 6000, Thermo Scientific, Thermo Fisher Scientific, Merelbeke, Belgium). Upon gas pressure measurement in the headspace, liquid samples were collected for subsequent analysis. Both the blank and probiotic treatment were tested for 6 different test subjects. For each test subject, multiple technical replicates were performed and harvested at either 0h (only blank), 3h, 6h, 9h or 24h.

Ileostomy samples were collected from healthy subjects after the participants signed an informed consent in which they donated their sample (procedure approved by Ethics Committee of the University Hospital Ghent; reference number BC-09977).

### Absolute microbial community analysis

Quantitative insights were obtained by correcting proportional data (%; shallow shotgun sequencing) with total cell counts for each sample (cells/mL; flow cytometry), resulting in estimated cell counts/mL.

DNA extraction was performed as previously described [27]. from the cell pellets obtained by centrifuging 1 mL culture for 1 min at 15.000 RPM. Details are depicted in the supplementary methods.

DNA libraries were prepared using the Nextera XT DNA Library Preparation Kit (Illumina) and IDT Unique Dual Indexes with total DNA input of 1 ng. Libraries were then sequenced on an Illumina Nextseq 2000 platform 2×150 bp. Unassembled sequencing reads were directly analyzed by CosmosID-HUB Microbiome Platform (CosmosID Inc., Germantown, MD) described elsewhere [28–31] for multi-kingdom microbiome analysis and profiling of antibiotic resistance and virulence genes and quantification of organisms’ relative abundance. Cleaned reads were assembled using metaSpades in default configuration [32]. Genes were predicted using Prodigal in metagenomics mode and subsequently functions were assigned by EggnogMapper [33, 34]. Details regarding library preparation, sequencing and bioinformatics analysis are depicted in the supplementary methods.For total cell count analysis, liquid samples were diluted in anaerobic phosphate-buffered saline, after which cells were stained with SYTO 16 at a final concentration of 1 μM and counted via a BD FACS Verse flow cytometer (BD, Erembodegem, Belgium). Data was analyzed using FlowJo, version 10.8.1.

### Extraction and LC-MS/MS

Metabolites for LC-MS/MS analysis were extracted from 350 μL culture supernatant using ethyl acetate. LC-MS/MS acquisition was performed using Shimadzu Nexera X2 UHPLC system, with attached PDA, coupled to Shimadzu 9030 QTOF mass spectrometer, equipped with a standard ESI source unit, in which a calibrant delivery system (CDS) is installed. All the samples were analyzed in positive and negative polarity, using data dependent acquisition mode. Full scan MS spectra (*m*/*z* 100–1700, scan rate 10 Hz, ID enabled) were followed by two data dependent MS/MS spectra (*m*/*z* 100–1700, scan rate 10 Hz, ID disabled) for the two most intense ions per scan. Details regarding the metabolite extraction and LC-MS/MS are depicted in the supplementary methods.

### ^1^H NMR spectroscopy and data processing

1 mL culture was centrifuged for 1 min at 21130 rcf, 250 μL of the supernatant was added to 400 μL NMR buffer (200 mM Na_2_HPO_4_, 44 mM NaH_2_PO_4_, 1 mM TSP, 3 mM NaN_3_ and 20% (v/v) D_2_O), centrifuged at 4 °C, 21130 rcf for 20 min and 550 μL was transferred to a 5 mm NMR tube.

All ^1^H-NMR spectra were recorded using a Bruker 600 MHz AVANCE II spectrometer equipped with a 5 mm triple resonance inverse cryoprobe and a z-gradient system. One-dimensional (1D) ^1^H-NMR spectra were recorded using the first increment of a NOESY pulse sequence with pre-saturation (γB_1_ = 50 Hz) for water suppression during a relaxation delay of 4 s and a mixing time of 10 ms. A 256 scans of 65,536 points covering 13,658 Hz were recorded and zero filled to 65,536 complex points prior to Fourier transformation, an exponential window function was applied with a line-broadening factor of 1.0 Hz. The spectra were phase and baseline corrected and referenced to the internal standard (TSP; δ 0.0 ppm), using the MestReNova software (v.12.0.0-20080, Mesterlab Research). The annotation of the bins was performed with the Chenomx Profiler software (Chenomx NMR Suite 8.6 and Chenomx 600 MHz, version 11) and the HMDB database 5.0 (http://www.hmdb.ca). Details regarding spectral imaging and data processing are depicted in the supplementary methods.

### Comparative metabolomics

Raw data obtained from the LC-MS analysis were converted to mzXML centroid files using Shimadzu LabSolutions Postrun Analysis. The files were imported into Mzmine 2.53 for data processing [35]. The resulting quantification tables were uploaded to MetaboAnalyst and subjected to RM two-way ANOVA [36]. The exported quantification table for GNPS and the MS2 spectra were uploaded to Feature Based Molecular Networking on the GNPS platform [37, 38] using the default settings. The resulting network was visualized in Cytoscape 3.4.0. Details regarding data processing are depicted in the supplementary methods.

### Statistical and network analysis

All analysis of variance (ANOVA) were performed using GraphPad Prism 7.0. When a comparison resulted in a statistically significant difference (P < 0.05), multiple comparisons testing was performed by controlling the false discovery rate (FDR) according to the Benjamini-Hochberg (BH) method (α < 0.05). The R package MixOmics was used for ordination and multivariate statistical analysis of the ^1^H-NMR spectra [39]. The dynamic profile comparison using Kendall’s τ correlation with the BH (α < 0.05) was performed using the R package psych. In CytoScape 3.9.1 the plugin CoNet [40] was used for network construction. The network properties were obtained using the network analyzer tool in cytoscape.

## Results

### Supplementation of a probiotic formula to the ileostoma microbiota alters the microbiota composition and function

To investigate the impact of probiotics supplementation on the small intestinal microbial community, healthy test subjects (n=6) collected their ileostoma effluent, which were used as a non-invasive access route to the otherwise inaccessible small intestine [24]. Ileostoma samples were inoculated in an ex vivo SIFR^®^ fermenter platform consisting of 54 bioreactors. Each ileostoma sample was incubated with or without a probiotic formula consisting of 9 probiotic species in separate bioreactors, at 37 °C with an initial pO_2_ of 15 mmHg, to simulate intraluminal oxygen levels of the proximal ileum [41] **(Figure 1A)**. The dynamic changes in the ileostoma microbiota composition upon probiotics supplementation were investigated using shallow shotgun metagenomic sequencing combined with flow cytometry on samples collected at different time points (0, 3, 6, 9, 24h) after incubation with or without the probiotic formula **(Figure 1A)**. Comparing the microbial richness, assessed by the Chao1 index (ordinary two-way ANOVA; F(1,49) = 2.1, *P* = 0.15 and F(4,49) = 0.15, *P* = 0.96) and the diversity as determined by Shannon’s H (ordinary two-way ANOVA; F(1,49) = 0.08, P = 0.77 and F(4,49) = 0.40, *P* = 0.81) and inverted Simpson’s index (ordinary two-way ANOVA; F(1,49) = 3.0, *P* = 0.088 and F(4,49) = 0.24, *P* = 0.91) did not result in significant differences between control and probiotics-supplemented samples **(Figure 1B)**. Though a notable decrease in the detected number of species after inoculation was observed irrespective of probiotics supplementation **(Figure 1B)**, the addition of probiotic cells to the ileostoma samples in the bioreactors decreased the absolute number of cells in the probiotics-supplemented samples compared to the control, except after 3h of growth **(Supplementary figure 1)**, indicating an altered microbiota composition.

**Figure 1:**
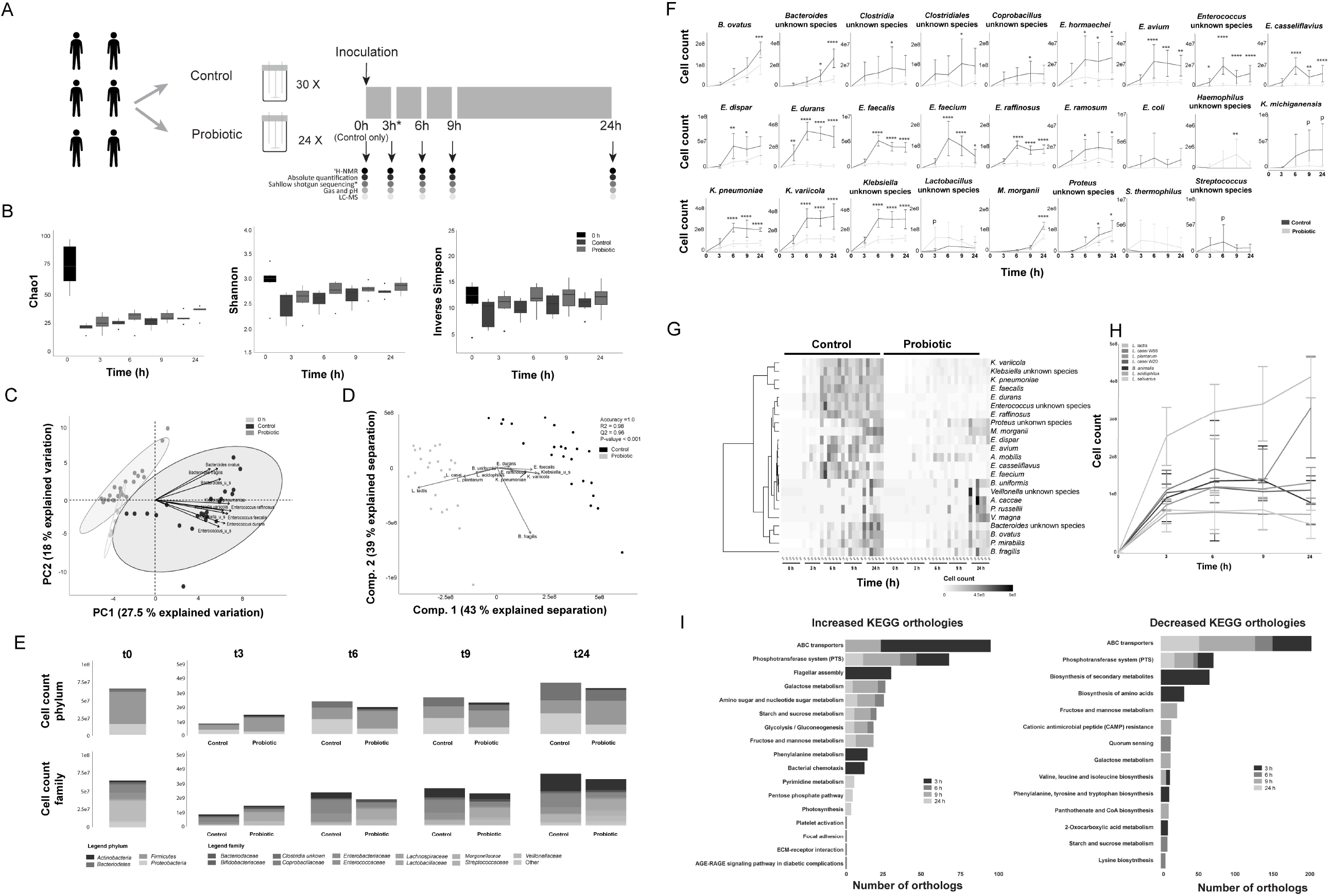
A probiotic formula reduces the cell counts of *Enterococcus* and *Klebsiella* species in compositional data obtained from ileostomy effluent. A) Schematic overview of the experimental procedure. Ileostomy effluent of 6 donors is collected and divided over 54 bioreactors. 24/54 bioreactors are supplemented with 10^7^ CFU/mL of the probiotic formula. The other 30/54 are controls without supplementation. The black arrows indicate sampling timepoints, where the contents of 6 control and 6 supplemented bioreactors are subjected to shallow shotgun metagenomic, flow cytometry, ^1^H-NMR, LC-MS and fermentation parameter analysis. The contents of 6 control bioreactors are subjected to the same analysis before the start of the experiment. The asterisk depicts the DNA sample obtained from donor 5 at timepoint 3h, which got destroyed during sample processing. Therefore this sample could not be used for the shallow shotgun sequencing analysis. B) The mean and standard deviation of the microbial richness as assessed by the Chao1 index and the microbial diversity as assessed by Shannon’s H and the inverse Simpson index. Differences are calculated using an ordinary two-way ANOVA. C) Principal component analysis of the relative abundance as obtained via shallow shotgun sequencing multiplied by the absolute cell count as obtained via flow cytometry. The ellipses represent the 95% confidence interval. The taxa contributing the most to the separation are indicated with arrows. D) Partial least square – discriminant analyses of the relative abundance as obtained via shallow shotgun sequencing multiplied by the absolute cell count as obtained via flow cytometry. The taxa contributing the most to the separation are depicted with arrows. The accuracy, R2 and Q2 values are statistical parameters which estimate the predictive ability of the model and are calculated via cross validation. The P-value is the results of a permutation test performed with the separation distance statistic and 1000 permutations. All parameters are calculated using MetaboAnalyst 5.0. E) Depiction of the average cell counts timepoint at the phylum and family levels as calculated by multiplying the relative abundance data obtained from the shallow shotgun sequencing and the absolute cell counts as obtained via flow cytometry. F) Dynamic cell count profiles of significantly different taxa between the control and probiotics-supplemented ileostoma microbiotas according to the factor condition as calculated by an ordinary two-way ANOVA with multiple comparisons testing by controlling the false discovery rate according to the Benjamini-Hochberg procedure; *: q < 0.05, **: q < 0.01, ***: q < 0.001, ****: q < 0.0001, P: P < 0.05, q > 0.05. G) Heatmap showing the significantly different taxa between the control and probiotics-supplemented ileostoma microbiota according to the factor time as calculated by an ordinary two-way ANOVA with multiple comparisons testing by controlling the false discovery rate according to the Benjamini-Hochberg procedure. H) Dynamic cell count profiles of 7/9 species present in the probiotic formula. *Bifidobacterium animalis* W51 and W52 could not be distinguished and *Bifidobacterium bifidum* W23 could not be identified in the shallow shotgun sequencing data. I) Differences in KEGG orthologs within the microbiota of the control vs probiotics-supplemented community. The differences are obtained by comparing the orthologs of the control and probiotics-supplemented community per timepoint using a linear modeling approach.

To investigate the changes in the absolute abundance of the ileostoma microbiota, the relative shallow shotgun data was multiplied by the absolute cell count data per sample **(Supplementary data 1)**. To only investigate probiotics affected species, the species only detected in the samples obtained before fermentation (0h) **(Supplementary data 1)** were excluded. Principal component analysis (PCA) of the remaining cell count data showed distinct clustering of the probiotics-supplemented, and control samples, mainly due to variations in the absolute abundance of multiple species of *Bacteroides, Klebsiella* and *Enterococcus* **(Figure 1C)**. This was further supported by partial least square – discriminant analysis (PLS-DA) (*P* < 0.001) **(Figure 1D)**. Next, compositional changes over time were compared to identify which taxa were affected by the supplementation of the probiotic formula. Focusing on the phylum level, the abundance of Firmicutes (ordinary one-way ANOVA; F(4,24) = 20.5, *P* < 0.0001), Proteobacteria (ordinary one-way ANOVA; F(4,24) = 3.8, *P* = 0.016) and Bacteroidetes (ordinary one-way ANOVA; F(4,24) = 11.4, *P* < 0.0001) were significantly altered throughout the course of the experiment, particularly at 3h and 24h of the fermentation in the control samples **(Figure 1E, supplementary figure 2)**. Although the abundance of Firmicutes (ordinary one-way ANOVA; F(4,25) = 5.6, *P* = 0.002), Actinobacteria (ordinary one-way ANOVA; F(4,25) = 6.1, *P* = 0.001) and Bacteroidetes (ordinary one-way ANOVA; F(4,25) = 13.0, *P* < 0.0001) were significantly altered throughout the course of the experiment, only the abundance of Actinobacteria was altered at 3h of the experiment in the probiotics-supplemented samples. The changes in abundance of Firmicutes and Bacteroidetes were most notable after 24h. In contrast to the control samples, the levels of Proteobacteria did not change throughout the experiment in the probiotics-supplemented samples (ordinary one-way ANOVA; F(4,25) = 2.5, *P* = 0.07) **(Figure 1E, supplementary figure 2)**. During and after termination of the experiment, no Actinobacteria were detected in the control samples, while we detected Actinobacteria in the probiotics-supplemented community throughout and after termination of the experiment **(Figure 1E, supplementary figure 2)**. The difference in the levels of Actinobacteria and the differences in the levels of Firmicutes between control and probiotics-supplemented samples are likely due to the composition of the probiotic formula, which contains seven Firmicutes and two Actinobacteria species. On the family level, the abundance of *Enterobacteriaceae* (ordinary one-way ANOVA; F(4,24) = 6.3, *P* = 0.001), *Bacteroidaceae* (ordinary one-way ANOVA; F(4,24) = 12.9, *P* < 0.0001) and *Enterococcaceae* (ordinary one-way ANOVA; F(4,24) = 13.1, *P* < 0.0001) were altered throughout the experiment with the most prominent changes after 3h and 24h of growth in the control samples, while the abundance of *Enterobacteriaceae* (ordinary one-way ANOVA; F(4,25) = 2.6, *P* = 0.06) did not significantly change throughout the experiment in the probiotics-supplemented samples. The abundance of *Bacteroidaceae* (ordinary one-way ANOVA; F(4,25) = 14.6, *P* < 0.0001) and *Enterococcaceae* (ordinary one-way ANOVA; F(4,25) = 58.6, *P* < 0.0001) were significantly altered throughout the experiment in the presence of the probiotics. **(Figure 1E, supplementary figure 2)**. To investigate how the microbiota composition changed on the species level, the differences in absolute levels per species were compared between the control and probiotics-supplemented samples utilizing an ordinary two-way ANOVA approach. Only 3/26 significantly different taxa (*Streptococcus thermophilus*, an unidentified *Haemophilus* and *Lactobacillus* species) were higher in the probiotics-supplemented samples **(Figure 1F)**, while 23/26 taxa, mainly *Enterobacteriaceae, Bacteroidaceaea* and *Enterococcaceae* were significantly lower **(Figure 1F)**, even though most species increased in number over time in both the control and probiotics-supplemented samples **(Figure 1G)**, confirming the ordination analysis **(Figure 1C and D)**. Investigation of the growth of the probiotic species showed only an increase in cell counts of *L. casei* W56 (ordinary one-way ANOVA; F(3,20) = 6.5, *P* = 0.003) and *L. lactis* W19 (ordinary one-way ANOVA; F(3,20) = 4.2, *P* = 0.018) over time **(Figure 1H)**.

To determine the alterations in the metabolic pathways associated with the addition of the probiotic formula to the ileostoma samples, metagenomic functional profiling on KEGG orthologs (KO) was performed. Throughout the experiment, 17 KO had increased and 14 KO had decreased **(Figure 1I)**. The KO: flagellar assembly, bacterial chemotaxis and phenylalanine metabolism were more abundant, whereas biosynthesis of secondary metabolites, biosynthesis of amino acids, phenylalanine, tyrosine and tryptophan biosynthesis and, 2-oxocarboxylic acid metabolism were less abundant after 3h of incubation in the probiotics-supplemented samples. At every sampling point the biggest changes (increase or decrease) were observed in KO describing phosphotransferase system (PTS) and ABC transporters **(Figure 1I)**, indicating that, throughout the course of the experiment, the bacteria which were able to take up and utilize available metabolites could survive. Altogether, the addition of the probiotic formula to the adult human small intestinal ileostoma microbiota caused a dynamic alteration in the function and composition of the community mainly due to a significant decreased representation of multiple species of *Bacteroides, Klebsiella* and *Enterococcus* over time.

### Altered metabolic activity of the ileostoma microbiota in response to the probiotic formula supplementation

Alterations in the microbiota composition and function are likely accompanied by alterations in the metabolic environment [42]. To investigate the dynamic changes in the metabolic environment of the ileostoma samples, fermentation parameters were obtained, and the samples were subjected to ^1^H-NMR and LC-MS analysis **(Figure 1A)**. The fermentation parameters, pH, and gas production decreased over time in both conditions. Of note, the reduction in the measured parameters was significantly lower in the probiotics-supplemented samples compared to the control samples at all time-points, hinting towards an altered metabolism **(Figure 2A, 2B)**. To further investigate the metabolic changes associated with the addition of the probiotic formula, we applied ordination analysis of binned and processed ^1^H-NMR spectra. The analysis revealed a time-dependent clustering of the spectra, which was driven by acetate (bin 387 and 388), propionate (bin 214), lactate (bin 268 and 270), acetone (bin 450), ethanol (bin 240) and an unidentified compound (bin 744, 745 and 746) **(Figure 2C)**. PLS-DA showed a distinct separation between the control and probiotics-supplemented samples, largely driven by differences in lactate (bin 268, 269, 270 and 271) and ethanol (bin 823, 825, 826, 828 and 830) **(Figure 2D)**. As early as 3h of community growth, a separation between the control and probiotics-supplemented ileostoma samples was detected, with lactate (bin 268 and 270) as the biggest contributor to the separation **(Supplementary figure 3A)**. No donor specific separation was observed **(Supplementary figure 3B)**. Furthermore, repeated measure (RM) two-way

**Figure 2:**
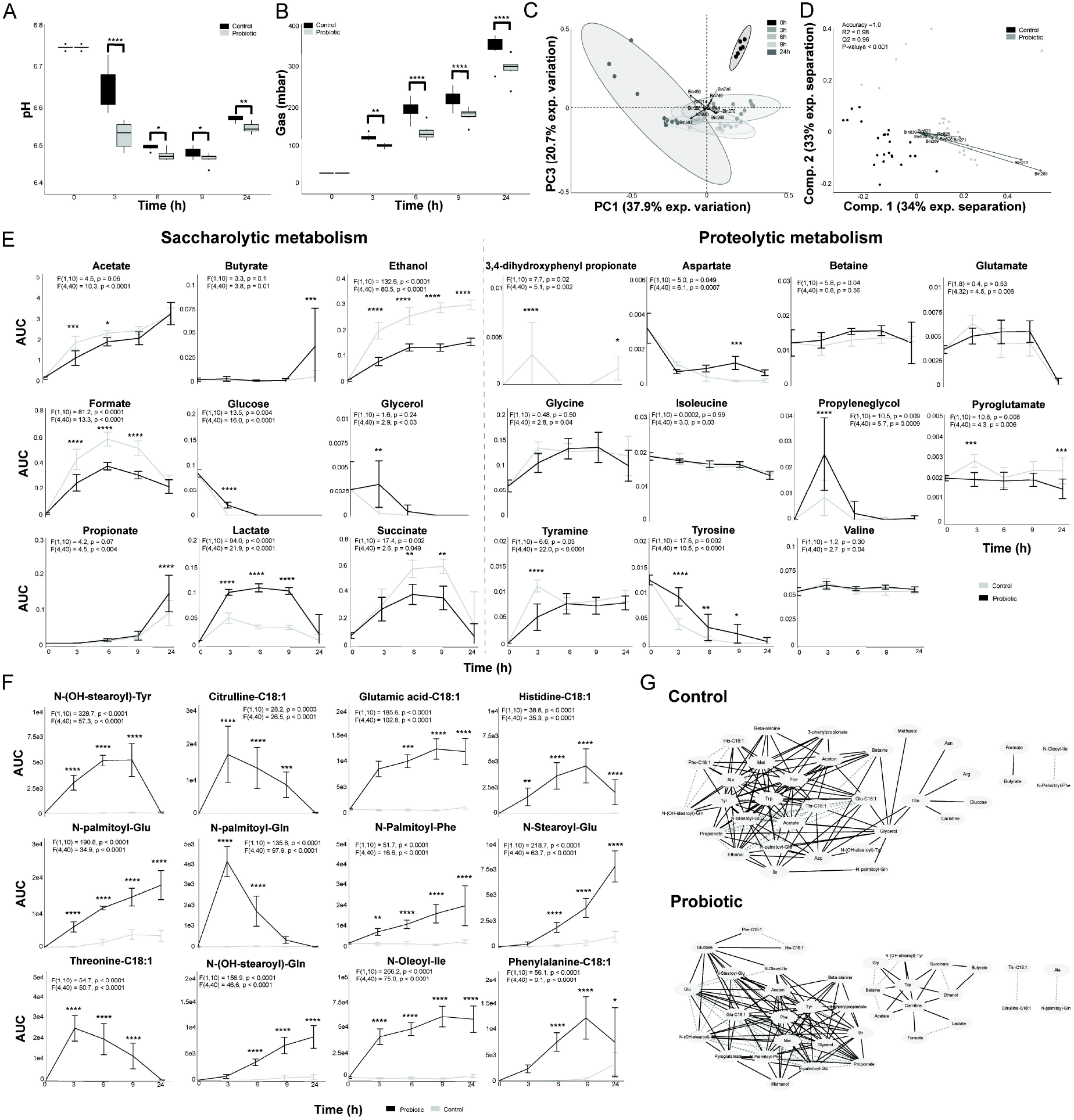
A probiotic formula alters the metabolic environment when supplemented to ileostomy effluent. A) The mean and standard deviation of the pH measured at each timepoint. B) The mean and standard deviation of the gas production at each timepoint. For panel B and C: Significance is obtained via a repeated measure two-way ANOVA with multiple comparison testing by controlling the false discovery rate according to the Benjamini-Hochberg procedure; *: q < 0.05, **: q < 0.01, ***: q < 0.001, ****: q < 0.0001. C) Principal component analysis performed with the binned spectra as input. The ellipses represent the 95 % confidence interval. The 10 bins contributing the most to the separation are indicated with arrows and belong to *propionate (bin 214), acetate (bin 387 and 388), lactate (bin 268 and 270), acetone (bin 450), ethanol (bin 240) and an unidentified compound (bin 744, 745 and 746)*. D) Partial least square – discriminant analyses performed with the binned spectra as input. The 9 bins contributing the most to the separation are indicated with arrows and belong *to lactate (bin 268, 269, 270 and 271) and ethanol (bin 823, 825, 826, 828 and 830)*. The accuracy, R2 and Q2 values are statistical parameters which estimate the predictive ability of the model and are calculated via cross validation. The *P*-value is the results of a permutation test performed with the separation distance statistic and 1000 permutations. All parameters are calculated using MetaboAnalyst 5.0. E) Dynamic profiles of the significantly different saccharolytic and proteolytic metabolites as obtained by measuring the area under the curve of a representative peak per metabolite. The error bars indicate the standard deviation. Significant differences are obtained via repeated measure two-way ANOVA. The F statistic and P value for the factor condition and for the interaction between time and condition are indicated for each metabolite. The result of multiple comparison testing by controlling the false discovery rate according to the Benjamini-Hochberg procedure is indicated with asterisks; *: q < 0.05, **: q < 0.01, ***: q < 0.001, ****: q< 0.0001. F) Dynamic profiles of the significantly different and identified metabolites as measured the area under the curve obtained by LC-MS. The error bars indicate the standard deviation. Significant differences are obtained via repeated measure two-way ANOVA. The F statistic and P value for the factor condition and for the interaction between time and condition are indicated for each metabolite. The result of multiple comparison testing by controlling the false discovery rate according to the Benjamini-Hochberg procedure is indicated with asterisks; *: q < 0.05, **: q < 0.01, ***: q < 0.001, ****: q < 0.0001. G) Dynamic correlation based networks of all metabolites measured in the control and probiotics-supplemented community. Nodes represent metabolites. Edges represents correlations between the two connected nodes and are obtained when 2/3 methods give a positive results; Kendall’s −0.8 > τ > 0.8, Spearman’s −0.8 > ρ > 0.8 and Brown’s randomization method with 1000 iterations has a Benjamini-Hochberg corrected P-value < 0.05. The calculation of the edges is performed using the Cytoscape plugin CoNet. Black solid edges represent negative correlations, grey dashed edges represents positive correlations.

ANOVA identified higher levels of saccharolytic metabolites such as butyrate (24h), propionate (24h) and lactate (3, 6 and 9h) and lower levels of acetate (3 and 6h) in the probiotics-supplemented ileostoma samples compared to the control samples. In contrast, the levels of succinate (6 and 9h), ethanol (3, 6, 9 and 24h) and formate (3, 6 and 9h) were reduced in the probiotics-supplemented samples **(Figure 2E)**. Among the altered proteolytic metabolic profiles, RM two-way ANOVA detected lower levels of tyrosine and higher levels of tyramine after 3h of growth in the probiotics-supplemented samples. However, the levels of tyramine in the probiotics-supplemented samples were not different compared to the control after 6, 9 and 24h of growth. Similarly, the levels of the saccharolytic metabolites; glucose, glycerol and propyleneglycol were only higher after 3h of growth in the probiotics-supplemented samples **(Figure 2E)**. Notably, the proteolytic metabolite, 3,4-dihydroxyphenylpropionic acid was the only metabolite exclusively detected in the control samples **(Figure 2E)**. Apart from the metabolites altered upon probiotics supplementation, the levels of several dynamically altered metabolites were similar between control and probiotics-supplemented samples **(Supplementary figure 4)**.

Next to the ^1^H-NMR, LC-MS was employed to investigate the effect of probiotics supplementation on metabolites outside the detection limits of ^1^H-NMR. Analogous to the ^1^H-NMR data, RM two-way ANOVA on the LC-MS data identified 64 and 98 features in negative and positive mode respectively as significantly different between the control and probiotics-supplemented samples **(Supplementary data 2)**. The analysis of the interaction between time and probiotics supplementation revealed 62 and 66 features in negative and positive mode respectively as significantly different between control and probiotics-supplemented samples **(Supplementary data 2)**. Out of the detected features, three metabolites (*N*-Oleoyl-isoleucine, citrulline-C18:1 and glutamate-C18:1) could be identified (identification level 2 [43]) using the global natural products social molecular networking web platform [37, 38]. Additionally, nine more features could be dereplicated based on their high-resolution MS and MS/MS spectra as threonine-C18:1, histidine-C18:1, phenylalanine-C18:1, *N*-palmitoyl-glutamate, *N*-palmitoyl-glutamine, *N*-palmitoyl-phenylalanine, *N*-stearoyl-glutamic acid, *N*-hydroxystearoyl-tyrosine, and *N*-hydroxystearoyl-glutamine (level of identification is 3 [43]) **(Supplementary figure 5, 6 and 7)**. Importantly, all the identified metabolites were *N*-acyl-amino acid conjugates **(Figure 2F)**.

To investigate the global impact of probiotics supplementation on the microbial community and to infer the coordinate behavior of metabolites, whole network topology features such as the clustering coefficient (indicative of resistance to perturbations) and network density (indicative of nutrient utilization) [44] were compared. To do so, dynamic correlation-based networks were constructed using the Cytoscape plugin CoNet [40], whereby the nodes represent the 31 identified ^1^H-NMR and 12 identified LC-MS metabolites, the edges represent correlations between the dynamic metabolic profiles per condition as obtained via ^1^H-NMR and LC-MS data. The control network consisted of 34/43 nodes with 132 edges (clustering coefficient = 0.413, network density = 0.299) **(Figure 2G)**. While probiotics supplementation only slightly increased the number of nodes (35/43) and edges (135), the clustering coefficient (0.658) and network density (0.529) increased substantially **(Figure 2G)**, indicating that the ileostoma microbiota together with the probiotic species is more resistant to external perturbations and is better suited to utilize the metabolites available in the environment compared to the ileostoma microbiota without the supplemented probiotics. To investigate which nodes in the networks had the largest impact on the global organization, node parameters such as the node degree, which is the number of edges per node, and the node betweenness centrality, which is a measure of the number of shortest paths between any two nodes that pass through the node [45, 46], were compared **(Supplementary data 3)**. The largest increase in node degree due to probiotics supplementation was observed for *N*-palmitoyl-phenylalanine (13), glucose (10) and glutamate (9). The largest decrease in node degree due to probiotics supplementation was observed for alanine (11), acetate (11) and threonine-C18:1 (13). Additionally, the probiotics supplementation altered the node betweenness centrality, with the largest increase for carnitine (0.5) and glucose (0.2) and the largest decrease for glycerol (0.25) and glutamate (0.22) **(Supplementary data 3)**. Together, these results indicate that the altered metabolic activity in the probiotics-supplemented ileostoma microbiota generated a dynamic network which makes the community more resistant to external perturbations and more capable to utilize nutrients.

### Increased lactate and N-acyl amino acid production by the probiotic formula correlates with reduced growth of Enterobacteriaceae, Enterococcaceae and ethanol production

To investigate whether the changes observed in the community metabolism upon probiotics supplementation can be explained by the changes in microbiota composition, we employed Kendall’s τ correlation analysis **(Figure 3, supplementary data 4)**. The probiotic species correlated positively with each other and with the increase of lactate in the metabolic environment, while other lactic acid bacteria such as *Streptococcaceae* and *Enterococcaceae* [47] correlated negatively with lactate as well as with the probiotic species. Acetate and propionate correlated positively with *Bacteroides, Morganella* and *Proteus* species, which were significantly reduced in the probiotics-supplemented samples. Propionate and butyrate correlated positively with *Anaerostipes caccae* and butyrate also showed a positive correlation with an unknown *Anaerostipes* species and *Peptostreptococcus russellii*, although none of these species showed significant difference in their abundance between the control and probiotics-supplemented samples **(Figure 2E)**. Ethanol correlated negatively with the probiotic species, but correlated positively with multiple *Enterococcaceae* and *Enterobacteriaceae* species, implying a direct relationship. In contrast, the N-acyl amino acids were negatively correlated with the *Enterococcaceae*, but positively correlated with each other and the probiotic species. 3,4-dihydroxyphenylpropionic acid, a metabolite produced from the microbial metabolism of caffeic acid, was not detected in the probiotics-supplemented samples, while it positively correlated with tyramine, an unknown *Morganella* and *Streptococcus* species. Glutamate, which had a large increase in node degree, but a large decrease in node betweenness centrality in the probiotics-supplemented samples **(Figure 2G)**, correlated negatively with propionate, an unknown *Morganella* species and *A. caccae* and positively with multiple metabolites including lactate and tyrosine, inferring a link between the change in pH and the network topology. The metabolites glucose, glycerol, phenylalanine and propylene glycol, which differ in node degree, betweenness centrality or levels after 3h of growth **(Figure 2E, 2G)**, correlated positively with each other and with tyrosine, tryptophan and methionine, but negatively correlated with multiple *Bacteroides, Morganella* and *Klebsiella* species and ethanol, implying involvement in the response of the ileostoma microbiota to the supplementation with the probiotics. Together, the correlations indicate that the compositional changes of the adult human small intestinal stoma microbiota **(Figure 1F)** after the addition of the probiotic formula were due to alterations in the metabolic environment **(Figure 2E, 2F)**.

## Discussion

This study investigated the impact of the supplementation of a 9-species probiotic formula on the dynamic metabolic interaction network of the microbiota of ileostoma samples obtained from healthy subjects. The network topology features showed that the supplementation of the probiotic formula increased the resistance to external perturbations and the nutrient utilization in the adult human small intestinal stoma microbiota **(Figure 2G)**. This action appears to be mediated via two mechanisms; pH mediated competitive exclusion and alteration of the metabolic interactions.

PH mediated competitive exclusion within the microbial community occurred via increased production of lactate and the subsequent increase in acidity, which is the most prevalent probiotic mechanism of action [48–50]. This mechanism has been linked to eradication of several pathogens, including *C. difficile, Escherichi coli* and *Klebsiella pneumonia*, increased production of short chain fatty acids [51, 52] and reduced production of succinate and formate [53, 54]. In line with the previous reports, our data showed an exclusion of *Enterococcaceae, Bacteriodaceae* and *Enterobacteriaceae* from the ileostoma community **(Figure 1F)**, higher levels of butyrate and propionate after 24h of the addition of the probiotics and lower levels of succinate and formate over the course of the experiment **(Figure 2E)**.

The increased acidity of the microbial environment resulted in activation of bacterial acid stress mechanisms to maintain their intracellular pH [55]. *Enterococcaceae*, one of the most dominant families in the small intestine **(Figure 1E)** [24], tolerate acid stress via activation of a tyrosine decarboxylase [56], an enzyme which converts tyrosine to tyramine. The present data infers higher tyrosine decarboxylase activity in the control ileostoma samples because the levels of tyramine and tyrosine significantly decreased and increased, respectively, compared to the samples supplemented with the probiotic formula **(Figure 2A, 2E)**, which is opposite to our hypothesis. However, these results could be explained by the significant decrease in the absolute abundance of *Enterococcus* species upon the addition of probiotics **(Figure 1F)**. This scenario is further supported by the negative correlations detected between multiple Enterococcus species and the probiotic species as well as the lower abundance of *Enterococcus* species upon probiotics supplementation **(Figure 3A)**. Analogously, Fernandez et al. found a reduction of *Enterococcus durans growth* with a lower pH [57].

**Figure 3:**
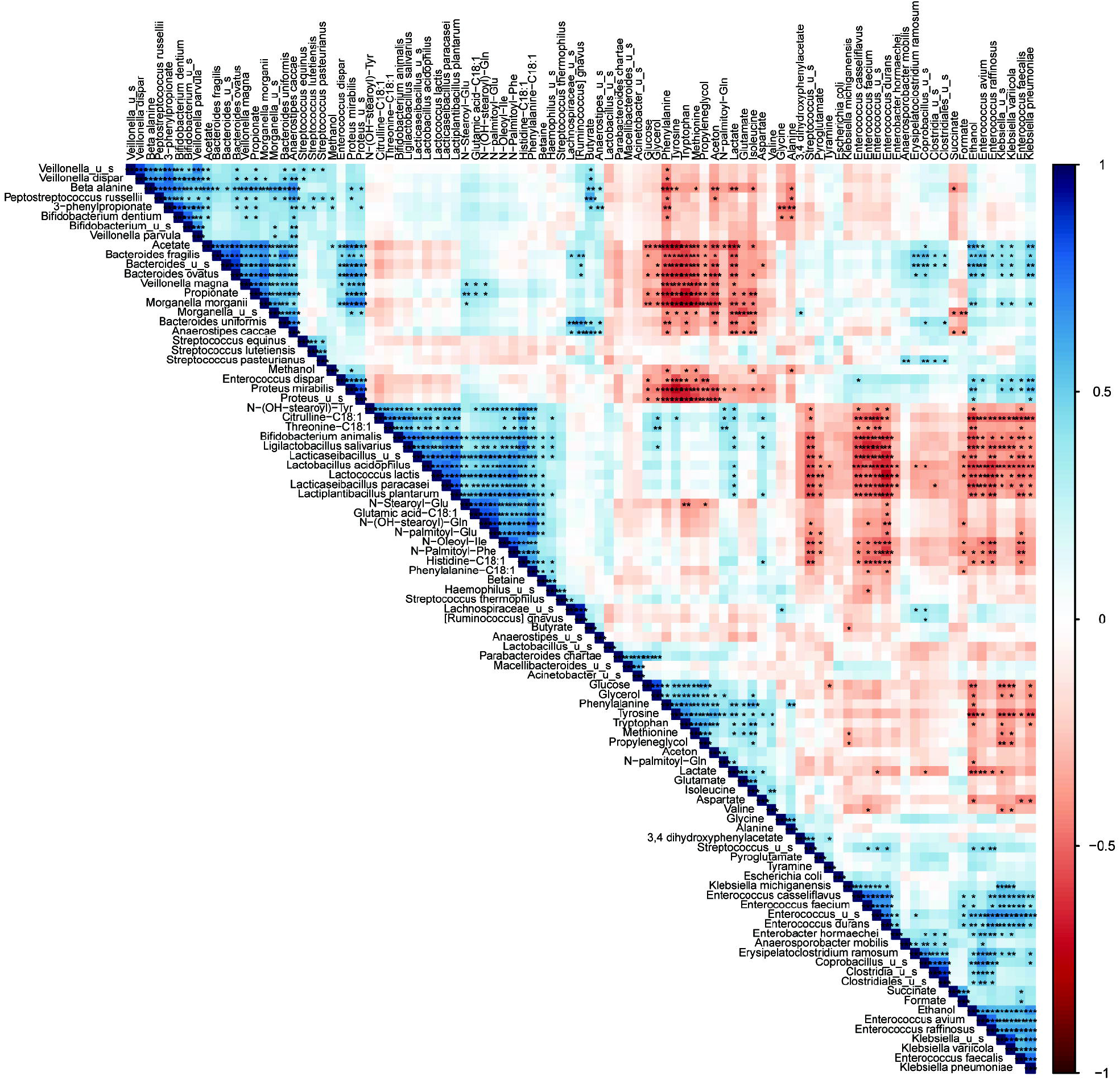
Dynamic correlations within the microbiota show a different dynamic metabolic fingerprint of the community with and without probiotic supplementation. Kendall’s τ correlation matrix of all metabolites and bacterial species present. Positive correlations are indicated in blue and negative correlations are indicated in red according the legend on the right. Significance is calculated by controlling the false discovery rate according to the Benjamini-Hochberg correction method and indicated with asterisks; *: q < 0.05, **: q < 0.01, ***: q < 0.001.

**Figure 4:**
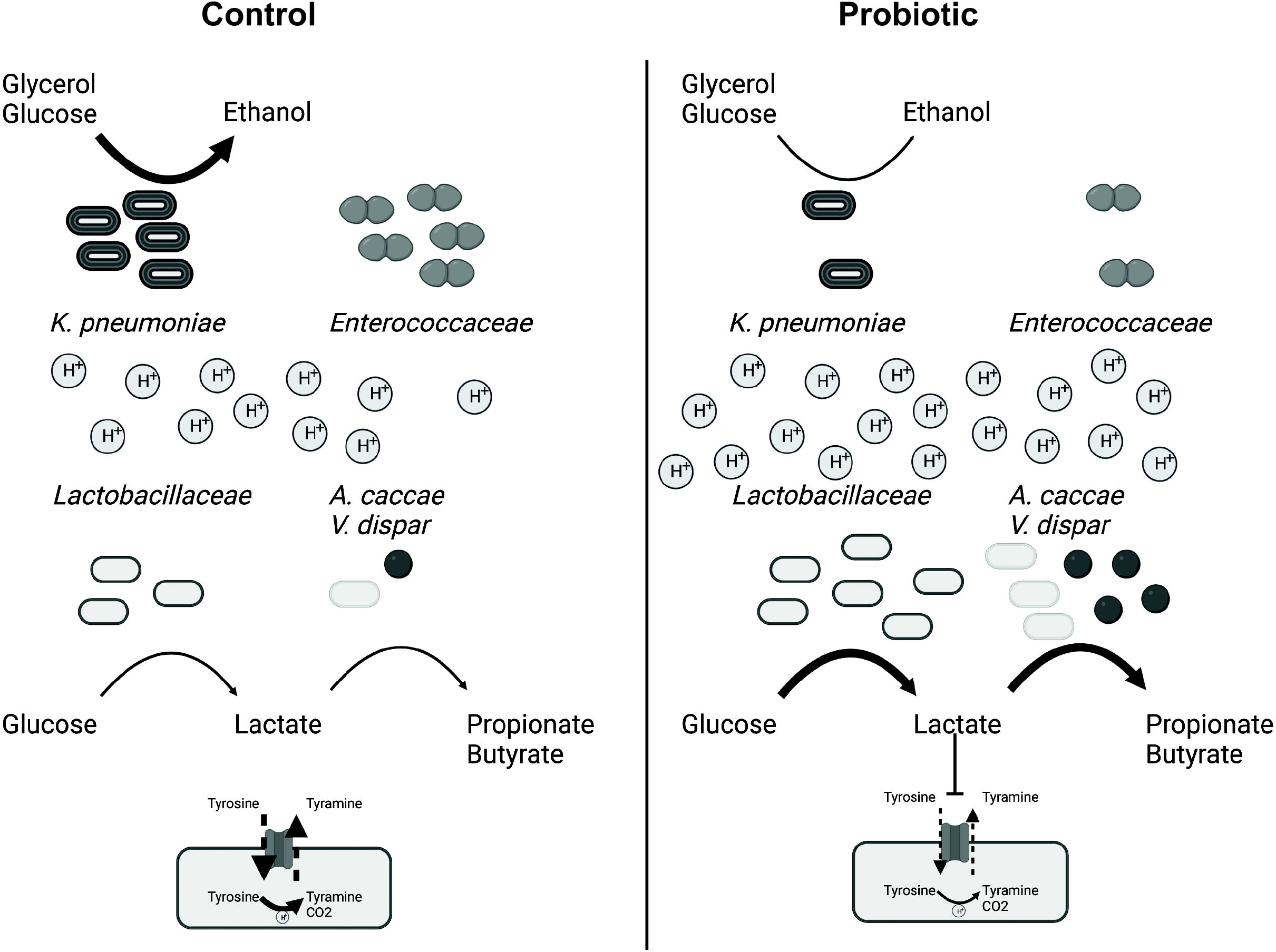
the probiotics supplementation to the ileostoma microbiota alters the metabolic environment and microbiota composition in a pH dependent manner. Supplementation of the probiotic formula increase the level of *Lactobacillaceae*, which reduces the pH of the metabolic environment by increasing the production of lactate, which also increases the production of propionate and butyrate by *Anaestipes cacae* and *Veillonella dispar*. Furthermore, the cells counts of *Enterococcaceae* are reduced due to lactate since lactate inhibits the tyrosine decarboxylase of *Enterococcoceae*, which is their main pH stress regulator. Additionally, the levels of *Klebsielle pneumonia* are reduced. Subsequently reducing the conversion of glucose and glycerol into ethanol by *K. pneumoniae.*

The significant reduction in *Enterococcus* species could be related to the reduced levels of ethanol detected in the probiotic supplemented samples **(Figure 1F, 2E)**. Indeed, our correlation analysis shows negative correlations between ethanol and the probiotic species, and positive correlations between ethanol and multiple *Enterococcus* species **(Figure 3A)**, in agreement with previous observations showing a positive association between Enterococcus levels and increased ethanol levels and pH in the small intestine, which promotes alcoholic liver disease [58]. The reduced levels of ethanol observed in the present study may be related to the lower abundance of *K. pneumoniae* upon probiotics supplementation **(Figure 1F)**. *K. pneumoniae*, which has been detected in the normal healthy human microbiota [24, 59, 60], can produce high levels of ethanol, endogenously in humans, from glucose and glycerol via the 2,3-butanediol fermentation pathway, and this has been linked to nonalcoholic fatty liver disease [61, 62]. However, the growth of K. pneumoniae is reduced at acidic pH [63, 64]. This is in agreement with our correlation analysis, which showed a positive correlation between ethanol and *Klebsiella* species, including *K. pneumoniae* and negative correlations between ethanol and glucose and glycerol, as well as a significant reduction of glycerol in the control community only after 3h of incubation **(Figure 1F, 2E, 3A)**, coinciding with the drop in the pH **(Figure 1A)**. The observed restricted growth of *Enterobacteriaceae* species in our data could explain the increase in absolute abundance of *Clostridiales* species (depicted in our data as *Anaerostipes*) and higher levels of butyrate production **(Figure 1G, 2E)**. In fact, it has been shown that interaction between *Lactobacillus* and *Clostridiales* species caused restriction in the growth of *Enterobacteriaceae* species and the authors hypothesized that *Lactobacillus* supplementation could promote the *Clostridiales* growth [65]. Together, the probiotic formula used in this study may have elicited the increased resistance to perturbations and the metabolic usage in the stoma community by lowering the pH, reducing ethanol production via reduction of *K. pneumonia* growth, consequently lowering *Enterococcus* growth. Subsequently, the increased lactate levels, most likely, resulted in higher production of propionate and butyrate by *Anearostipes* and *Veillonella* species **(Figure 4)**.

Although the higher bacterial cell count had a direct impact on the levels of the produced metabolites, the interaction of specific bacterial strains within the probiotic formula with the surrounding microbial community may be equally important in determining the effect of the supplementation of probiotics. Among the 9 bacterial strains that constitute the probiotic formula, our data revealed an increase in the cell count of only 2/9 species **(Figure 1H)**, inferring that the metabolic interaction with the surrounding bacteria rather than the growth of the probiotic strains was the determinant factor of the measured effects of the probiotics. To understand these metabolic interactions, we constructed metabolic networks based on the dynamic profile of the identified metabolites. The increase in network density and the clustering coefficient of the metabolic networks **(Figure 2G)** further support the importance of the metabolic interactions within the community. Our observations are in agreement with Jeong et al. who compared the network diameter among metabolic networks of individual organisms and compared these networks with non-biological networks. The network diameter is the largest number of edges between two nodes in the network and is a measure of connectivity within the network, comparable to the network density. The authors showed that the addition of a new node to a non-biological network increased the network diameter, thereby reducing the connectivity. In contrast, the addition of nodes to biological networks did not alter the network diameter, implying a large resistance to external perturbations and tolerance to node removal [66], which is supported by the low number of essential genes in bacteria [67, 68]. Closer inspection of the changed network showed that N-palmitoyl-phenylalanine and threonine-C18:1 were among the metabolites with the largest differences in network impact. Moreover, these two metabolites positively correlated with the probiotic species and N-palmitoyl-phenylalanine positively correlated with the other identified N-acyl-amino acids **(Figure 3)**. Additionally, all *N*-acyl-amino acids were all produced in higher amounts in the probiotics-supplemented samples **(Figure 2F)**, however, this increase does not necessarily mean they are directly produced by the probiotic strains but due to the changes in the microbiota composition that followed the supplementation. Indeed, gut bacteria, notably *Clostridia*, can produce fatty acid conjugates [69], and these metabolites can be used by the gut microbiota as signaling molecules, and have been found to act as antimicrobials[69, 70]. Although the precise biological role of the N-acyl-amino acids in the gut is largely unknown [71], these molecules have been categorized, together with the endocannabinoid system, as the endocannabinoidome [72] and alterations in the endocannabinoidome have been implicated in a variety of gut related disorders [73, 74]. Further research is needed to elucidate the precise role of the N-acyl-amino acids within the microbiota and between microbiota and host.

Overall, the present study shows that combining metabolic network construction with compositional analysis of adult human small intestinal stoma microbiota supplemented with a mixture of probiotics revealed a dynamic increase in the resistance of the community to perturbations and nutrient utilization. These alterations were the consequences of a pH related competitive exclusion of *Enterobacteriaceae* and *Enterococcaceae*. The outcome may advance our understanding of the mechanisms underlying the beneficial host effects associated with the supplementation of probiotics and help improve the current or identify novel microbiota-targeted therapies, particularly in the small intestine being a pivotal target for the probiotic activity.

## Author Contributions

JJ, SEA: Conceptualization, investigation, writing-original draft, and editing; JJ, MS, PvA, NuT: data analysis; JJ, ACC, NuT, SsES, PvA: methodology; SvH, GpvW: Conceptualization, writing review; SEA: funding acquisition. All authors have read and agreed to the published version of the manuscript.

## Data availability

The raw LC-MS/MS data was deposited in the MASSIVE database, accession ID: MSV000091160 The shallow shotgun sequencing data were deposited under BioProject number PRJNA928311

## References

1. Faith JJ, Guruge JL, Charbonneau M, Subramanian S, Seedorf H, Goodman AL, et al. The long-term stability of the human gut microbiota. Science (1979) 2013; 341.

2. Fassarella M, Blaak EE, Penders J, Nauta A, Smidt H, Zoetendal EG. Gut microbiome stability and resilience: elucidating the response to perturbations in order to modulate gut health. Gut 2021; 70: 595–605.

3. Kern L, Abdeen SK, Kolodziejczyk AA, Elinav E. Commensal inter-bacterial interactions shaping the microbiota. Curr Opin Microbiol 2021; 63: 158–171.

4. Weiss AS, Burrichter AG, Chakravarthy A, Raj D, von Strempel A, Meng C, et al. In vitro interaction network of a synthetic gut bacterial community. The ISME Journal 2021 2021; 1–15.

5. Layeghifard M, Hwang DM, Guttman DS. Disentangling Interactions in the Microbiome: A Network Perspective. Trends Microbiol 2017; 25: 217–228.

6. Barabási AL, Gulbahce N, Loscalzo J. Network medicine: a network-based approach to human disease. Nature Reviews Genetics 2011 12:1 2010; 12: 56– 68.

7. Sung J, Kim S, Cabatbat JJT, Jang S, Jin YS, Jung GY, et al. Global metabolic interaction network of the human gut microbiota for context-specific community-scale analysis. Nature Communications 2017 8:1 2017; 8: 1–12.

8. Greenblum S, Turnbaugh PJ, Borenstein E. Metagenomic systems biology of the human gut microbiome reveals topological shifts associated with obesity and inflammatory bowel disease. Proc Natl Acad Sci U S A 2012; 109: 594–599.

9. Rosario D, Bidkhori G, Lee S, Bedarf J, Hildebrand F, le Chatelier E, et al. Systematic analysis of gut microbiome reveals the role of bacterial folate and homocysteine metabolism in Parkinson’s disease. Cell Rep 2021; 34: 108807.

10. Dohlman AB, Shen X. Mapping the microbial interactome: Statistical and experimental approaches for microbiome network inference. Exp Biol Med 2019; 244: 445.

11. Steinway SN, Biggs MB, Loughran TP, Papin JA, Albert R. Inference of Network Dynamics and Metabolic Interactions in the Gut Microbiome. PLoS Comput Biol 2015; 11: e1004338.

12. Hill C, Guarner F, Reid G, Gibson GR, Merenstein DJ, Pot B, et al. The International Scientific Association for Probiotics and Prebiotics consensus statement on the scope and appropriate use of the term probiotic. Nature Reviews Gastroenterology & Hepatology 2014 11:8 2014; 11: 506–514.

13. Zucko J, Starcevic A, Diminic J, Oros D, Mortazavian AM, Putnik P. Probiotic – friend or foe? Curr Opin Food Sci 2020; 32: 45–49.

14. Ritchie ML, Romanuk TN. A Meta-Analysis of Probiotic Efficacy for Gastrointestinal Diseases. PLoS One 2012; 7: e34938.

15. Puvanasundram P, Chong CM, Sabri S, Yusoff MSM, Lim KC, Karim M. Efficacy of Single and Multi-Strain Probiotics on In Vitro Strain Compatibility, Pathogen Inhibition, Biofilm Formation Capability, and Stress Tolerance. Biology (Basel) 2022; 11: 1644.

16. Bagga D, Reichert JL, Koschutnig K, Aigner CS, Holzer P, Koskinen K, et al. Probiotics drive gut microbiome triggering emotional brain signatures. Gut Microbes 2018; 9: 486–496.

17. Moser AM, Spindelboeck W, Halwachs B, Strohmaier H, Kump P, Gorkiewicz G, et al. Effects of an oral synbiotic on the gastrointestinal immune system and microbiota in patients with diarrhea-predominant irritable bowel syndrome. Eur J Nutr 2019; 58: 2767–2778.

18. Wilms E, Gerritsen J, Smidt H, Besseling-Van Der Van Vaart I, Rijkers GT, Fuentes ARG, et al. Effects of supplementation of the synbiotic Ecologic® 825/FOS P6 on intestinal barrier function in healthy humans: A randomized controlled trial. PLoS One 2016; 11.

19. Martinez-Guryn K, Hubert N, Frazier K, Urlass S, Musch MW, Ojeda P, et al. Small intestine microbiota regulate host digestive and absorptive adaptive responses to dietary lipids. Cell Host Microbe 2018; 23: 458.

20. Rios-Morales M, van Trijp MPH, Rösch C, An R, Boer T, Gerding A, et al. A toolbox for the comprehensive analysis of small volume human intestinal samples that can be used with gastrointestinal sampling capsules. Scientific Reports 2021 11:1 2021; 11: 1–14.

21. Kastl AJ, Terry NA, Wu GD, Albenberg LG. The Structure and Function of the Human Small Intestinal Microbiota: Current Understanding and Future Directions. Cell Mol Gastroenterol Hepatol 2020; 9: 33–45.

22. Matsuzawa H, Munakata S, Kawai M, Sugimoto K, Kamiyama H, Takahashi M, et al. Analysis of ileostomy stool samples reveals dysbiosis in patients with high-output stomas. Biosci Microbiota Food Health 2021; 40: 135.

23. Colom J, Freitas D, Simon A, Brodkorb A, Buckley M, Deaton J, et al. Presence and Germination of the Probiotic Bacillus subtilis DE111® in the Human Small Intestinal Tract: A Randomized, Crossover, Double-Blind, and Placebo-Controlled Study. Front Microbiol 2021; 12: 2189.

24. Yilmaz B, Fuhrer T, Morgenthaler D, Krupka N, Wang D, Spari D, et al. Plasticity of the adult human small intestinal stoma microbiota. Cell Host Microbe 2022; 30: 1773-1787.e6.

25. van den Abbeele P, Deyaert S, Thabuis C, Perreau C, Bajic D, Wintergerst E, et al. Bridging Preclinical and Clinical Gut Microbiota Research Using the Ex Vivo SIFR Technology. Frontiers in Microbiology (under review) 2023.

26. de Vos WM, Tilg H, van Hul M, Cani PD. Gut microbiome and health: mechanistic insights. Gut 2022; 71: 1020–1032.

27. Yu Z, Morrison M. Improved extraction of PCR-quality community DNA from digesta and fecal samples. Biotechniques 2004; 36: 808–812.

28. Lax S, Smith DP, Hampton-Marcell J, Owens SM, Handley KM, Scott NM, et al. Longitudinal analysis of microbial interaction between humans and the indoor environment. Science (1979) 2014; 345: 1048–1052.

29. Hasan NA, Young BA, Minard-Smith AT, Saeed K, Li H, Heizer EM, et al. Microbial community profiling of human saliva using shotgun metagenomic sequencing. PLoS One 2014; 9.

30. Ponnusamy D, Kozlova E v, Sha J, Erova TE, Azar SR, Fitts EC, et al. Cross-talk among flesh-eating Aeromonas hydrophila strains in mixed infection leading to necrotizing fasciitis.

31. Ottesen A, Ramachandran P, Reed E, White JR, Hasan N, Subramanian P, et al. Enrichment dynamics of Listeria monocytogenes and the associated microbiome from naturally contaminated ice cream linked to a listeriosis outbreak. BMC Microbiol 2016; 16.

32. Nurk S, Meleshko D, Korobeynikov A, Pevzner PA. MetaSPAdes: A new versatile metagenomic assembler. Genome Res 2017; 27: 824–834.

33. Cantalapiedra CP, Hern□andez-Plaza A, Letunic I, Bork P, Huerta-Cepas J. eggNOG-mapper v2: Functional Annotation, Orthology Assignments, and Domain Prediction at the Metagenomic Scale. Mol Biol Evol 2021; 38: 5825–5829.

34. Huerta-Cepas J, Szklarczyk D, Heller D, Hernández-Plaza A, Forslund SK, Cook H, et al. eggNOG 5.0: a hierarchical, functionally and phylogenetically annotated orthology resource based on 5090 organisms and 2502 viruses. Nucleic Acids Res 2019; 47: D309–D314.

35. Pluskal T, Castillo S, Villar-Briones A, Orešič M. MZmine 2: Modular framework for processing, visualizing, and analyzing mass spectrometry-based molecular profile data. BMC Bioinformatics 2010; 11: 1–11.

36. Xia J, Psychogios N, Young N, Wishart DS. MetaboAnalyst: a web server for metabolomic data analysis and interpretation. Nucleic Acids Res 2009; 37: W652.

37. Wang M, Carver JJ, Phelan V v., Sanchez LM, Garg N, Peng Y, et al. Sharing and community curation of mass spectrometry data with Global Natural Products Social Molecular Networking. Nature Biotechnology 2016 34:8 2016; 34: 828–837.

38. Nothias LF, Petras D, Schmid R, Dührkop K, Rainer J, Sarvepalli A, et al. Feature-based molecular networking in the GNPS analysis environment. Nature Methods 2020 17:9 2020; 17: 905–908.

39. Rohart F, Gautier B, Singh A, Lê Cao KA. mixOmics: An R package for ‘omics feature selection and multiple data integration. PLoS Comput Biol 2017; 13: e1005752.

40. Faust K, Raes J, Wilmes P, Heintz-Buschart A, Eiler A. CoNet app: inference of biological association networks using Cytoscape. F1000Research 2016 5:1519 2016; 5: 1519.

41. de Vos WM, Tilg H, van Hul M, Cani PD. Gut microbiome and health: mechanistic insights. Gut 2022; 71: 1020–1032.

42. Cao Y, Liu Y, Dong Q, Wang T, Niu C. Alterations in the gut microbiome and metabolic profile in rats acclimated to high environmental temperature. Microb Biotechnol 2022; 15: 276–288.

43. Sumner LW, Amberg A, Barrett D, Beale MH, Beger R, Daykin CA, et al. Proposed minimum reporting standards for chemical analysis: Chemical Analysis Working Group (CAWG) Metabolomics Standards Initiative (MSI). Metabolomics 2007; 3: 211–221.

44. Batushansky A, Matsuzaki S, Newhardt MF, West MS, Griffin TM, Humphries KM. GC-MS metabolic profiling reveals Fructose-2,6-bisphosphate regulates branched chain amino acid metabolism in the heart during fasting. Metabolomics 2019; 15: 18.

45. Vernocchi P, Gili T, Conte F, del Chierico F, Conta G, Miccheli A, et al. Network Analysis of Gut Microbiome and Metabolome to Discover Microbiota-Linked Biomarkers in Patients Affected by Non-Small Cell Lung Cancer. International Journal of Molecular Sciences 2020, Vol 21, Page 8730 2020; 21: 8730.

46. Tipton L, Müller CL, Kurtz ZD, Huang L, Kleerup E, Morris A, et al. Fungi stabilize connectivity in the lung and skin microbial ecosystems. Microbiome 2018; 6: 1–14.

47. Wang Y, Wu J, Lv M, Shao Z, Hungwe M, Wang J, et al. Metabolism Characteristics of Lactic Acid Bacteria and the Expanding Applications in Food Industry. Front Bioeng Biotechnol 2021; 9.

48. Mohan R, Koebnick C, Schildt J, Mueller M, Radke M, Blaut M. Effects of Bifidobacterium lactis Bb12 Supplementation on Body Weight, Fecal pH, Acetate, Lactate, Calprotectin, and IgA in Preterm Infants. Pediatric Research 2008 64:4 2008; 64: 418–422.

49. Kim HK, Rutten NBMM, Besseling-van der Vaart I, Niers LEM, Choi YH, Rijkers GT, et al. Probiotic supplementation influences faecal short chain fatty acids in infants at high risk for eczema. Benef Microbes 2015; 6: 783–790.

50. van Thu T, Foo HL, Loh TC, Bejo MH. Inhibitory activity and organic acid concentrations of metabolite combinations produced by various strains of Lactobacillus plantarum. Afr J Biotechnol 2013; 10: 1359–1363.

51. Sorbara MT, Dubin K, Littmann ER, Moody TU, Fontana E, Seok R, et al. Inhibiting antibiotic-resistant Enterobacteriaceae by microbiota-mediated intracellular acidification. J Exp Med 2019; 216: 84–98.

52. Kolling GL, Wu M, Warren CA, Durmaz E, Klaenhammer TR, Guerrant RL. Lactic acid production by Streptococcus thermophilus alters Clostridium difficile infection and in vitro Toxin A production. Gut Microbes 2012; 3: 523.

53. Ternes D, Tsenkova M, Pozdeev VI, Meyers M, Koncina E, Atatri S, et al. The gut microbial metabolite formate exacerbates colorectal cancer progression. Nature Metabolism 2022 4:4 2022; 4: 458–475.

54. Serena C, Ceperuelo-Mallafré V, Keiran N, Queipo-Ortuño MI, Bernal R, Gomez-Huelgas R, et al. Elevated circulating levels of succinate in human obesity are linked to specific gut microbiota. The ISME Journal 2018 12:7 2018; 12: 1642– 1657.

55. Guan N, Liu L. Microbial response to acid stress: mechanisms and applications. Appl Microbiol Biotechnol 2020; 104: 51–65.

56. Perez M, Calles-Enríquez M, Nes I, Martin MC, Fernandez M, Ladero V, et al. Tyramine biosynthesis is transcriptionally induced at low pH and improves the fitness of Enterococcus faecalis in acidic environments. Appl Microbiol Biotechnol 2015; 99: 3547–3558.

57. Fernández M, Linares DM, Rodríguez A, Alvarez MA. Factors affecting tyramine production in Enterococcus durans IPLA 655. Appl Microbiol Biotechnol 2007; 73: 1400–1406.

58. Llorente C, Jepsen P, Inamine T, Wang L, Bluemel S, Wang HJ, et al. Gastric acid suppression promotes alcoholic liver disease by inducing overgrowth of intestinal Enterococcus. Nature Communications 2017 8:1 2017; 8: 1–15.

59. Conlan S, Kong HH, Segre JA. Species-Level Analysis of DNA Sequence Data from the NIH Human Microbiome Project. PLoS One 2012; 7: e47075.

60. Zoetendal EG, Raes J, van den Bogert B, Arumugam M, Booijink CC, Troost FJ, et al. The human small intestinal microbiota is driven by rapid uptake and conversion of simple carbohydrates. The ISME Journal 2012 6:7 2012; 6: 1415– 1426.

61. Li NN, Li W, Feng JX, Zhang WW, Zhang R, Du SH, et al. High alcohol-producing Klebsiella pneumoniae causes fatty liver disease through 2,3-butanediol fermentation pathway in vivo. Gut Microbes 2021; 13.

62. Yuan J, Chen C, Cui J, Lu J, Yan C, Wei X, et al. Fatty Liver Disease Caused by High-Alcohol-Producing Klebsiella pneumoniae. Cell Metab 2019; 30: 675-688.e7.

63. Abbas SZ, Riaz M, Ramzan N, Zahid MT, Shakoori FR, Rafatullah M. Isolation and characterization of arsenic resistant bacteria from wastewater. Brazilian Journal of Microbiology 2014; 45: 1309.

64. Mitrea L, Vodnar DC. Klebsiella pneumoniae—A Useful Pathogenic Strain for Biotechnological Purposes: Diols Biosynthesis under Controlled and Uncontrolled pH Levels. Pathogens 2019; 8.

65. Djukovic A, Garzón MJ, Canlet C, Cabral V, Lalaoui R, García-Garcerá M, et al. Lactobacillus supports Clostridiales to restrict gut colonization by multidrug-resistant Enterobacteriaceae. Nature Communications 2022 13:1 2022; 13: 1– 18.

66. Jeong H, Tombor B, Albert R, Oltval ZN, Barabásl AL. The large-scale organization of metabolic networks. Nature 2000 407:6804 2000; 407: 651– 654.

67. Kobayashi K, Ehrlich SD, Albertini A, Amati G, Andersen KK, Arnaud M, et al. Essential Bacillus subtilis genes. Proc Natl Acad Sci U S A 2003; 100: 4678–4683.

68. Gerdes SY, Scholle MD, Campbell JW, Balázsi G, Ravasz E, Daugherty MD, et al. Experimental Determination and System Level Analysis of Essential Genes in Escherichia coli MG1655. J Bacteriol 2003; 185: 5673.

69. Chang FY, Siuti P, Laurent S, Williams T, Glassey E, Sailer AW, et al. Gut-inhabiting Clostridia build human GPCR ligands by conjugating neurotransmitters with diet- and human-derived fatty acids. Nature Microbiology 2021 6:6 2021; 6: 792–805.

70. Brady SF, Clardy J. Long-chain N-acyl amino acid antibiotics isolated from heterologously expressed environmental DNA [20]. J Am Chem Soc 2000; 122: 12903–12904.

71. Battista N, Bari M, Bisogno T. N-Acyl Amino Acids: Metabolism, Molecular Targets, and Role in Biological Processes. Biomolecules 2019, Vol 9, Page 822 2019; 9: 822.

72. di Marzo V. The endocannabinoidome as a substrate for noneuphoric phytocannabinoid action and gut microbiome dysfunction in neuropsychiatric disorders. Dialogues Clin Neurosci 2020; 22: 259.

73. Lian J, Casari I, Falasca M. Modulatory role of the endocannabinoidome in the pathophysiology of the gastrointestinal tract. Pharmacol Res 2022; 175: 106025.

74. Kim JT, Terrell SM, Li VL, Wei W, Fischer CR, Long JZ. Cooperative enzymatic control of N-ACYL amino acids by PM20D1 and FAAH. Elife 2020; 9.

