## Supplementary methods and figures for "Probiotics supplementation to adult human small intestinal stoma microbiota causes dynamic increase in the community resistance to perturbations and nutrient utilization"

*Fermentation experiment using SIFR® technology*

Individual bioreactors were processed in parallel in a bioreactor management device (Cryptobiotix, Ghent, Belgium). Each bioreactor contained 5 mL of nutritional medium (M0014, Cryptobiotix, Ghent, Belgium) - microbial inoculum blend with or without supplementation of the probiotic formula (at 10^7^ CFU probiotic/mL), then sealed individually, before being rendered anaerobic. At the start of the incubation, oxygen was introduced at pO_2_ = 15 mmHg, thus providing relevant intraluminal oxygen levels for the proximal ileum [1]. After preparation, bioreactors were incubated under continuous agitation (140 rpm) at 37 °C for 24h (MaxQ 6000, Thermo Scientific, Thermo Fisher Scientific, Merelbeke, Belgium). Upon gas pressure measurement in the headspace, liquid samples were collected for subsequent analysis. Both the blank and probiotic treatment were tested for 6 different test subjects. For each test subject, multiple technical replicates were performed and harvested at either 0h (only blank), 3h, 6h, 9h or 24h.

Ileostomy samples were collected from healthy subjects after the participants signed an informed consent in which they donated their sample (procedure approved by Ethics Committee of the University Hospital Ghent; reference number BC-09977).

*DNA extraction*

DNA was extracted from the cell pellets obtained by centrifuging 1 mL culture for 1 min at 21130 rcf. The supernatant was removed, the pellets were resuspended in 750 mL lysis buffer (500 mM NaCl, 50 mM Tris-HCl (pH 8.0), 50 mM EDTA, 4 % SDS) and transferred to a 2 mL screw cap tube containing 0.5 g, 0.1 mm zirconia beads and 4, 3 mm glass beads. The cells were disrupted with a mini-bead beater (3X 1 min, with 1 min intervals on ice) and incubated at 95 °C for 15 min. Cell debris was removed by centrifugation at 4 °C, 21130 rcf for 20 min. 500 µL of the supernatant was transferred to a clean tube. 200 µL 10 M ammonium acetate was added, and the samples were incubated for 10 min on ice. The samples were centrifuged at 4 °C, 21130 rcf for 10 min. 600 µL was transferred to a clean tube and 600 µL isopropanol was added. The samples were incubated overnight at -20 °C and afterwards centrifuged at 4 °C, 21130 rcf for 15 min to pellet the nucleic acids. The supernatant was removed and to wash the nucleic acid pellet 700 µL 70 % ethanol was added, the samples were mixed thoroughly and centrifuged at room temperature, 21130 rcf for 5 min. The pellets were dried by leaving the tubes open for 60 min at room temperature and were dissolved in 100 µL MilliQ.

*^1^H NMR spectroscopy and data processing*

1 mL culture was centrifuged for 1 min at 21130 rcf, 250 µL of the supernatant was added to 400 µL NMR buffer (200 mM Na­_2_HPO_4_, 44 mM NaH_2_PO_4_, 1 mM TSP, 3 mM NaN_3_ and 20% (v/v) D_2_O), centrifuged at 4 °C, 21130 rcf for 20 min and 550 µL was transferred to a 5 mm NMR tube.

All ^1^H-NMR spectra were recorded using a Bruker 600 MHz AVANCE II spectrometer equipped with a 5 mm triple resonance inverse cryoprobe and a z-gradient system. The temperature of the samples was controlled at 25 °C during measurement. Prior to data acquisition, tuning and matching of the probe head followed by shimming and proton pulse calibration were performed automatically for each sample. One-dimensional (1D) ^1^H-NMR spectra were recorded using the first increment of a NOESY pulse sequence with presaturation (γB_1_ = 50 Hz) for water suppression during a relaxation delay of 4 s and a mixing time of 10 ms. 256 scans of 65,536 points covering 13,658 Hz were recorded and zero filled to 65,536 complex points prior to Fourier transformation, an exponential window function was applied with a line-broadening factor of 1.0 Hz. The spectra were phase and baseline corrected and referenced to the internal standard (TSP; δ 0.0 ppm), using the MestReNova software (v.12.0.0-20080, Mesterlab Research). With the same software, spectral alignment was performed selecting manually areas and applying a linear filling method of the missing values, after the exclusion of the water signal and the surrounding empty area (4.50-5.30 ppm). Spectral binning followed from -0.50 to 9.00 ppm with an equal size binning step of 0.005 ppm. The noise removal was performed by averaging each integrated bin separately and removing the bins with an average below 4300. Before ordination analysis bins belonging to TSP (-0.50 – 0.72 ppm) and a pH sensitive area (7.850 – 8.190 ppm) were removed. The annotation of the bins was performed with the Chenomx Profiler software (Chenomx NMR Suite 8.6 and Chenomx 600 MHz, version 11) and the HMDB database 5.0 (<http://www.hmdb.ca>).

*Absolute microbial community analysis*

Quantitative insights were obtained by correcting proportional data (%; shallow shotgun sequencing) with total cell counts for each sample (cells/mL; flow cytometry), resulting in estimated cell counts/mL.

DNA extraction was performed as previously described [2]. from the cell pellets obtained by centrifuging 1 mL culture for 1 min at 15.000 RPM.

DNA libraries were prepared using the Nextera XT DNA Library Preparation Kit (Illumina) and IDT Unique Dual Indexes with total DNA input of 1 ng. Genomic DNA was fragmented using a proportional amount of Illumina Nextera XT fragmentation enzyme. Unique dual indexes were added to each sample followed by 12 cycles of PCR to construct libraries. DNA libraries were purified using AMpure magnetic Beads (Beckman Coulter) and eluted in QIAGEN EB buffer. DNA libraries were quantified using Qubit 4 fluorometer and Qubit™ dsDNA HS Assay Kit.
Libraries were then sequenced on an Illumina Nextseq 2000 platform 2x150 bp.

Unassembled sequencing reads were directly analyzed by CosmosID-HUB Microbiome Platform (CosmosID Inc., Germantown, MD) described elsewhere [3–6] for multi-kingdom microbiome analysis and profiling of antibiotic resistance and virulence genes and quantification of organisms' relative abundance. Briefly, the system utilizes curated genome databases and a high performance data-mining algorithm that rapidly disambiguates hundreds of millions of metagenomic sequence reads into the discrete microorganisms engendering the particular sequences. Similarly, the community resistome and virulome, the collection of antibiotic resistance and virulence associated genes in the microbiome, were also identified by querying the unassembled sequence reads against the CosmosID curated antibiotic resistance and virulence associated gene databases. Cleaned reads were assembled using metaSpades in default configuration [7]. Genes were predicted using Prodigal in metagenomics mode and subsequently functions were assigned by EggnogMapper [8, 9]. In order to estimate gene abundance, reads were mapped back to the assembly using Bowtie2 and coverage was assessed by employing Bedtools [10]. Coverage per gene was normalized by the total coverage of the sample and only genes with assigned KEGG orthologue (K-number) were retained. To assess differential abundance limma was used with ‘trend’ enabled [11]. Pathway enrichment was assessed by limma’s kegga and topKEGG methods. Taxonomy was assigned to all contigs by using CAT [12]. For total cell count analysis, liquid samples were diluted in anaerobic phosphate-buffered saline, after which cells were stained with SYTO 16 at a final concentration of 1 µM and counted via a BD FACS Verse flow cytometer (BD, Erembodegem, Belgium). Data was analyzed using FlowJo, version 10.8.1.

*Extraction and LC-MS/MS*

In a glass vial, 350 µL mL culture supernatant was mixed with 700 µL ethylacetate (EtOAc), and incubated overnight at 4 °C. Then, 350 µL of the upper organic layer was collected, and another 350 µL fresh EtOAc was added to the supernatant/EtOAc mixture and mixed. After a few hours of incubation, again 350 µL of the upper layer was collected and added to the first collection fraction. The solvent was evaporated from the collected extract under nitrogen flow. For LC-MS/MS analyses, the extracts were redissolved in 350 µL methanol.

LC-MS/MS acquisition was performed using Shimadzu Nexera X2 UHPLC system, with attached PDA, coupled to Shimadzu 9030 QTOF mass spectrometer, equipped with a standard ESI source unit, in which a calibrant delivery system (CDS) is installed. A total of 2 µL of the extracts were injected into a Waters Acquity HSS C_18_ column (1.8 μm, 100 Å, 2.1 × 100 mm). The column was maintained at 30 °C, and run at a flow rate of 0.5 mL/min, using 0.1% formic acid in H_2_O as solvent A, and 0.1% formic acid in acetonitrile as solvent B. A gradient was employed for chromatographic separation starting at 5% B for 1 min, then 5–85% B for 9 min, 85–100% B for 1 min, and finally held at 100% B for 3 min. The column was re-equilibrated to 5% B for 3 min before the next run was started. The LC flow was switched to the waste the first 0.5 min, then to the MS for 13.5 min, then back to the waste to the end of the run. The PDA acquisition was performed in the range 200–600 nm, at 4.2 Hz, with 1.2 nm slit width. The flow cell was maintained at 40 °C.

All the samples were analyzed in positive and negative polarity, using data dependent acquisition mode. Full scan MS spectra (*m/z* 100–1700, scan rate 10 Hz, ID enabled) were followed by two data dependent MS/MS spectra (*m/z* 100–1700, scan rate 10 Hz, ID disabled) for the two most intense ions per scan. The ions were selected when they reached an intensity threshold of 1500, isolated at the tuning file Q1 resolution, fragmented using collision induced dissociation (CID) with fixed collision energy (CE 20 eV), and excluded for 1 s before being re-selected for fragmentation. The parameters used for the ESI source were interface voltage 4 kV (positive polarity) or -3 kV (negative polarity), interface temperature 300 °C, nebulizing gas flow 3 L/min, and drying gas flow 10 L/min. The parameters used for the CDS probe were interface voltage 4.5 kV (positive polarity) or -3.5 kV (negative polarity), and nebulizing gas flow 1 L/min.

*Comparative metabolomics*

Raw data obtained from the LC-MS analysis was converted to mzXML centroid files using Shimadzu LabSolutions Postrun Analysis. The files were imported into Mzmine 2.53 for data processing [13]. Unless stated otherwise, m/z tolerance was set to 0.002 m/z or 15.0 ppm, RT tolerance was set to 0.1 min, noise level was set to 2.0E2 and the minimum absolute intensity was set to 5.0E2. Mass ion peaks were detected (positive polarity or negative polarity, mass detector: centroid) and their chromatograms were built using ADAP chromatogram builder (minimum group size in number of scans: 5; group intensity threshold: 2.0E2). The detected peaks were smoothed (filter width: 5), and the chromatograms were deconvoluted (algorithm: local minimum search; chromatographic threshold: 90 %; search minimum in RT range: 0.05; minimum relative height: 1 %; minimum ratio of peak top/edge: 2; peak duration: 0.03–3.00 min, MS/MS scan pairing (RT tolerance: 0.15 min, MS1 to MS2 precursor tolerance: 0.02 *m/z*)). The detected peaks were deisotoped (monotonic shape; maximum charge: 2; representative isotope: most intense). Peak lists from different extracts were aligned (weight for RT = weight for *m/z* = 20; compare isotopic pattern with a minimum score of 50%). Missing peaks detected in at least one of the samples were filled with the gap filling algorithm (Intensity tolerance: 50 %). Duplicate peaks were filtered (mode: New average, *m/z* tolerance: 0.002 *m/z* or 10 ppm, RT tolerance: 0.05 min). Artifacts caused by detector ringing were removed (*m/z* tolerance: 1.0 *m/z* or 1000.0 ppm, RT tolerance: 0.03 min). Only features with RT 0.5–14 min were kept. The aligned peaks were exported to a MetaboAnalyst file, as well as a GNPS file.

Features that were not present with an intensity higher than 3000 in at least 2 samples out of 6 from the same condition were removed. Additionally, all features that originate from the culture medium were removed by retaining only features with an average peak intensity of at least 20 times (positive polarity) or 10 times (negative polarity) greater in the bacterial extracts than in the culture medium extracts. The resulting peak lists were uploaded to MetaboAnalyst and subjected to RM two-way ANOVA [14]. The exported quantification table for GNPS and the MS2 spectra were uploaded to Feature Networking on the GNPS platform [15, 16] using the default settings for HR-MS/MS data. The settings regarding the negative mode can be found here: <https://gnps.ucsd.edu/ProteoSAFe/status.jsp?task=3b594ebd1dfd45f786fdd4591e963ee4>, the settings regarding the positive mode can be found here: <https://gnps.ucsd.edu/ProteoSAFe/status.jsp?task=e27430313b22469cb84f4d4091552734> The resulting network was visualized in Cytoscape 3.4.0.

**Supplementary Figures**

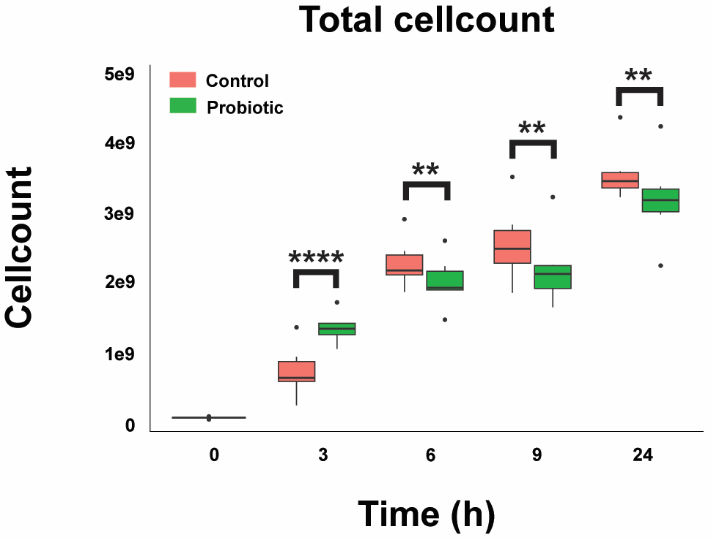

**Supplementary figure 1: Probiotic supplementation alters the total amount of cells when supplemented to ileostomy effluent**

The mean and standard deviation of the total cell count at each timepoint. Significance is obtained using a repeated measure two way ANOVA with multiple comparison testing by controlling the false discovery rate according to the Benjamini-Hochberg procedure; *: q < 0.05, **: q < 0.01, ***: q < 0.001, ****: q < 0.0001.

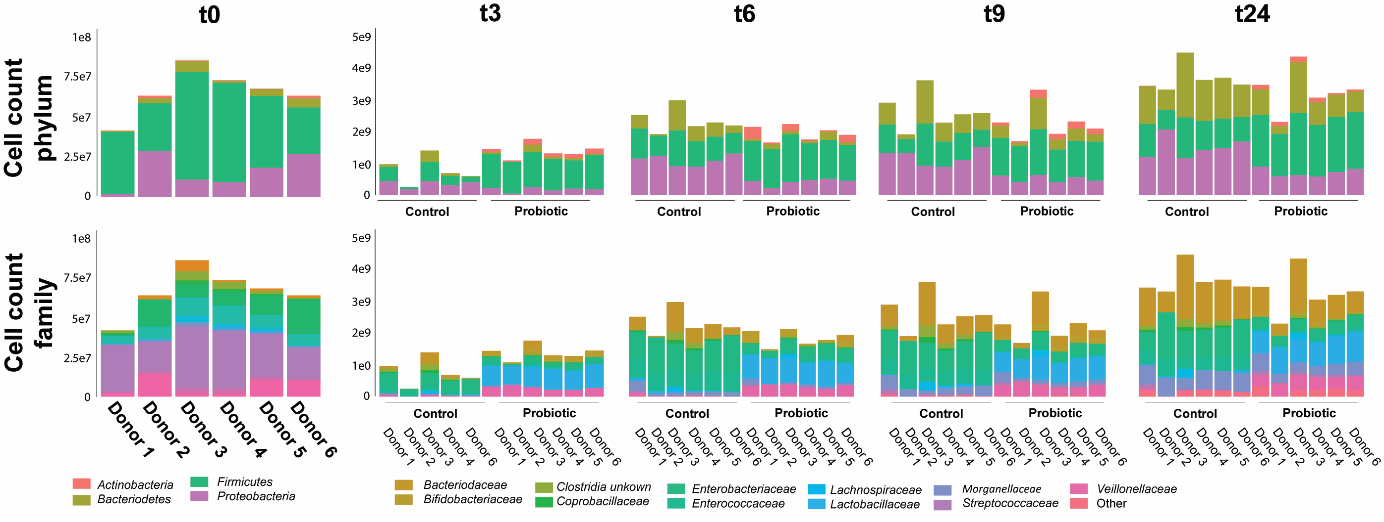

**Supplementary figure 2: cellcount per donor and timepoint at the phylum and family level**

Depiction of the cell counts per donor and timepoint at the phylum and family levels as calculated by multiplying the relative abundance data obtained from the shallow shotgun sequencing and the absolute cell counts as obtained via flow cytometry. During sample processing the DNA obtained from donor 5 at timepoint 3 got destroyed. Therefore this sample could not be used for the shallow shotgun sequencing analysis.

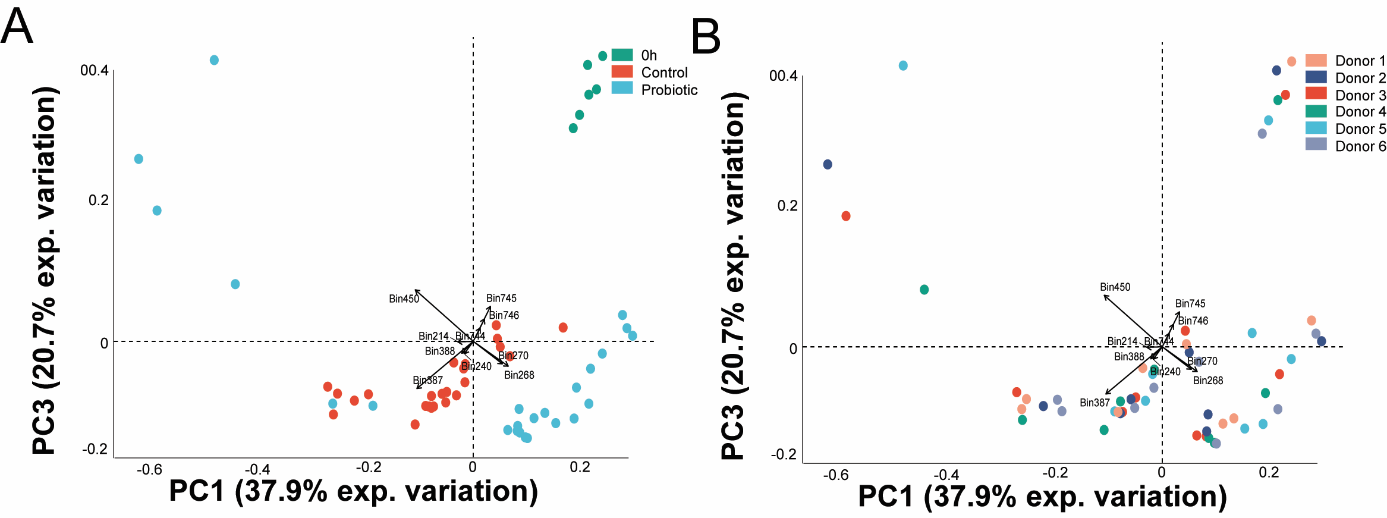

**Supplementary figure 3: The ^1^H-NMR spectra separate according to probiotic supplementation, not on donor**

Principal component analysis performed with the binned ^1^H-NMR spectra as input. The 10 bins contributing the most to the separation are indicated with arrows and belong to propionate (bin 214), acetate (bin 387 and 388), lactate (bin 268 and 270), acetone (bin 450), ethanol (bin 240) and an unidentified compound (bin 744, 745 and 746)*. A*) The samples are colored according to the samples obtained at 0h, the control and probiotic supplementation stoma samples. B) The samples are colored according to the donor.

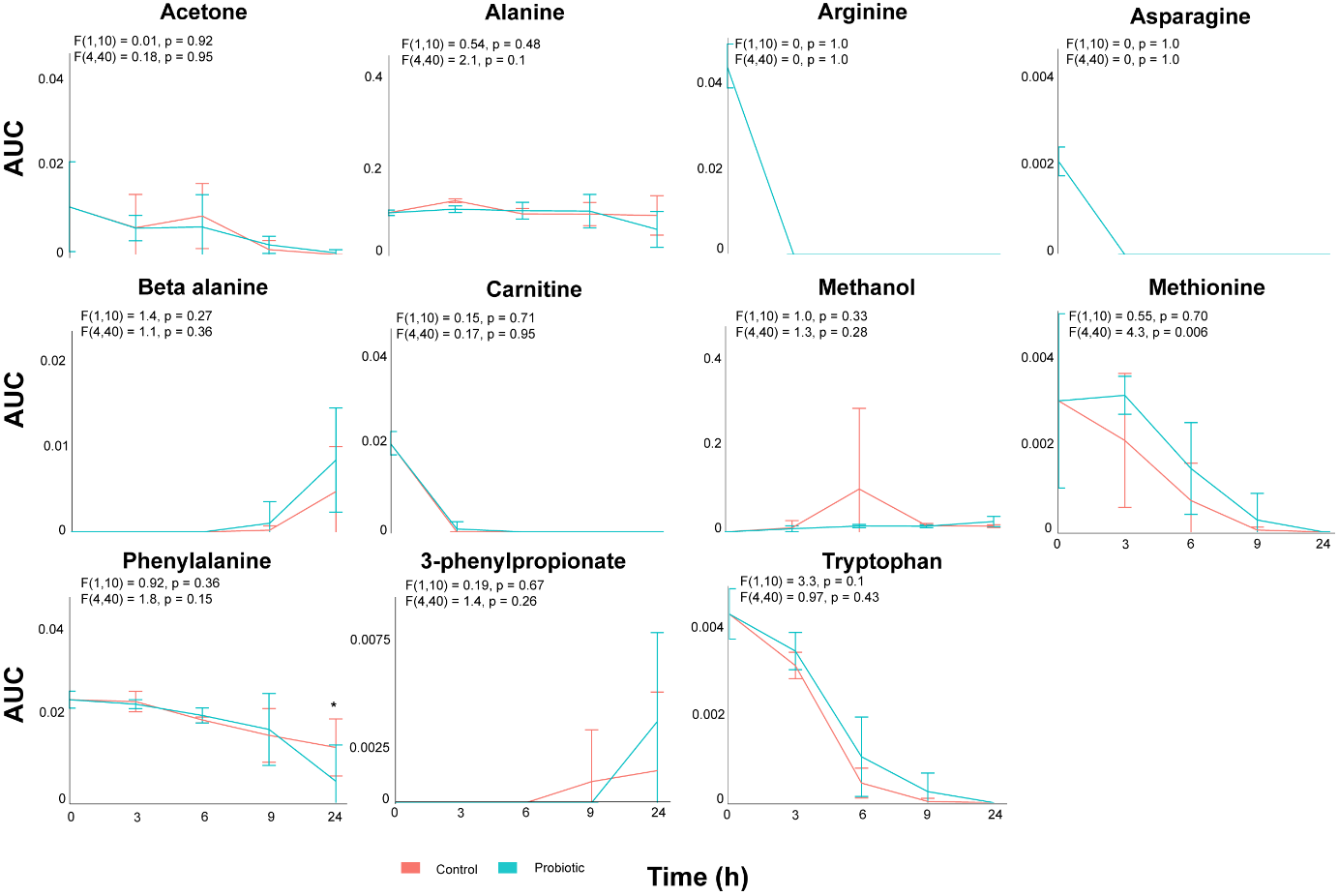

**Supplementary figure 4: Similar dynamic metabolic profiles of the stoma samples with and without probiotic supplementation**

Non significantly different dynamic metabolic profiles as obtained by measuring the area under the curve of a representative peak per metabolite. The error bars indicate the standard deviation per timepoint. Differences are investigated via repeated measure two way ANOVA. The F statistic and P value for the factor condition and for the interaction between time and condition are indicated for each metabolite. The result of multiple comparison testing by controlling the false discovery rate according to the Benjamini-Hochberg procedure is indicated with asterisks; *: q < 0.05.

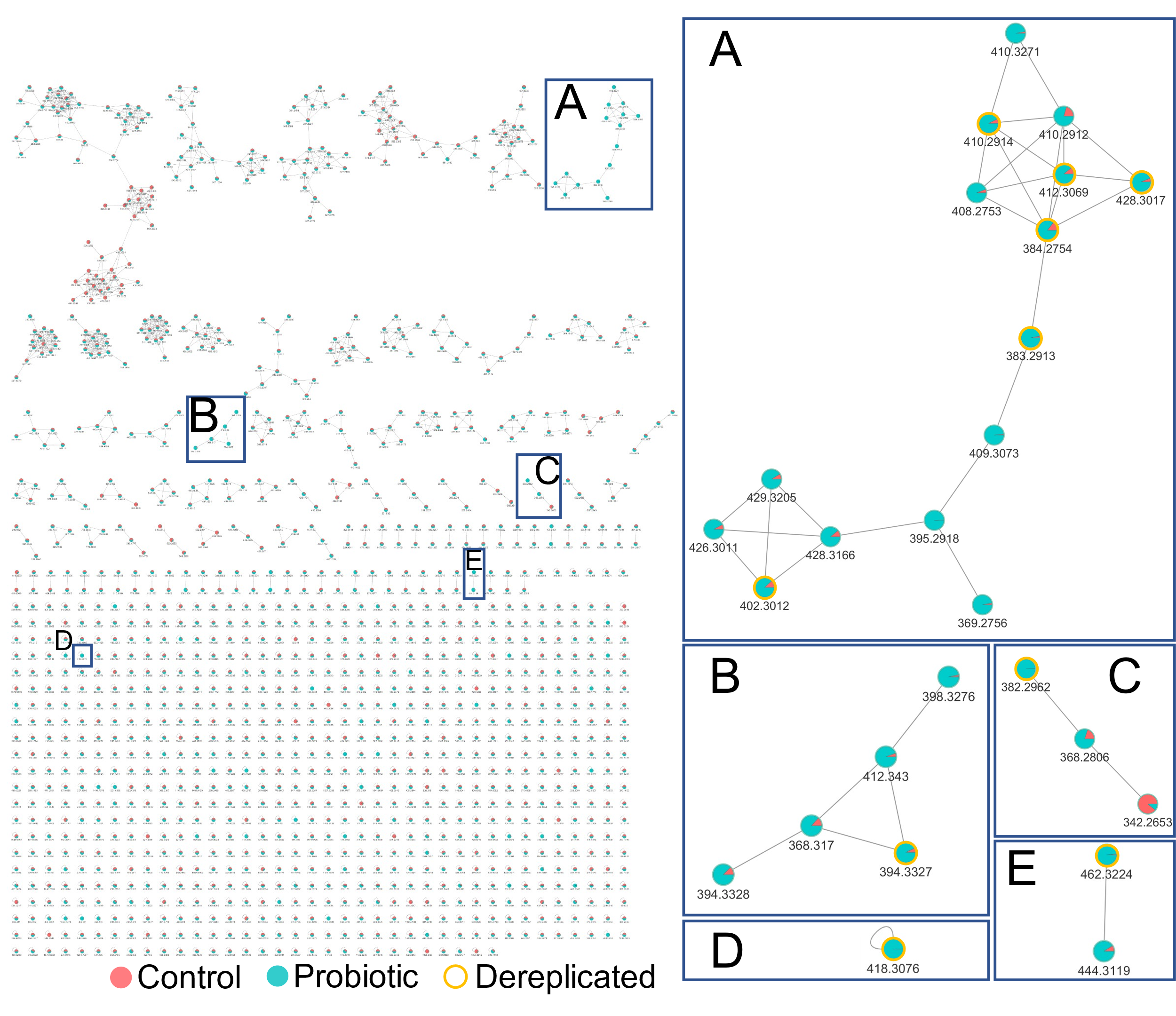

**Supplementary figure 5:** **Feature-based molecular network (FBMN) of the ions detected in the negative polarity (LC-MS/MS)**

Every node of the network represents a mass feature with a specific retention time, and is labelled with the *m/z* of the parent ion. The mass features are clustered together based on similar fragmentation patterns in their MS/MS spectrum. A pie chart is mapped to the nodes that shows the abundance of each feature in the control (salmon) and probiotic-supplemented (blue) samples. Dereplicated features are highlighted in yellow in the enlargements (A-E). **A** Glu-C18:1 (*m/z* 410.2914), *N*-stearoyl-Glu (*m/z* 412.3069), *N*-hydroxystearoyl-Gln (*m/z* 428.3017), *N*-palmitoyl-Glu (*m/z* 384.2754), *N*-palmitoyl-Gln (*m/z* 383.2913), *N*-palmitoyl-Phe (*m/z* 402.3012) **B** *N*-oleoyl-Ile (*m/z* 394.3327) **C** Thr-C18:1 (*m/z* 382.2962) **D** His-C18:1 (*m/z* 418.3076) **E** *N*-hydroxystearoyl-Tyr (*m/z* 462.3224). The FBMN was illustrated using Cytoscape.

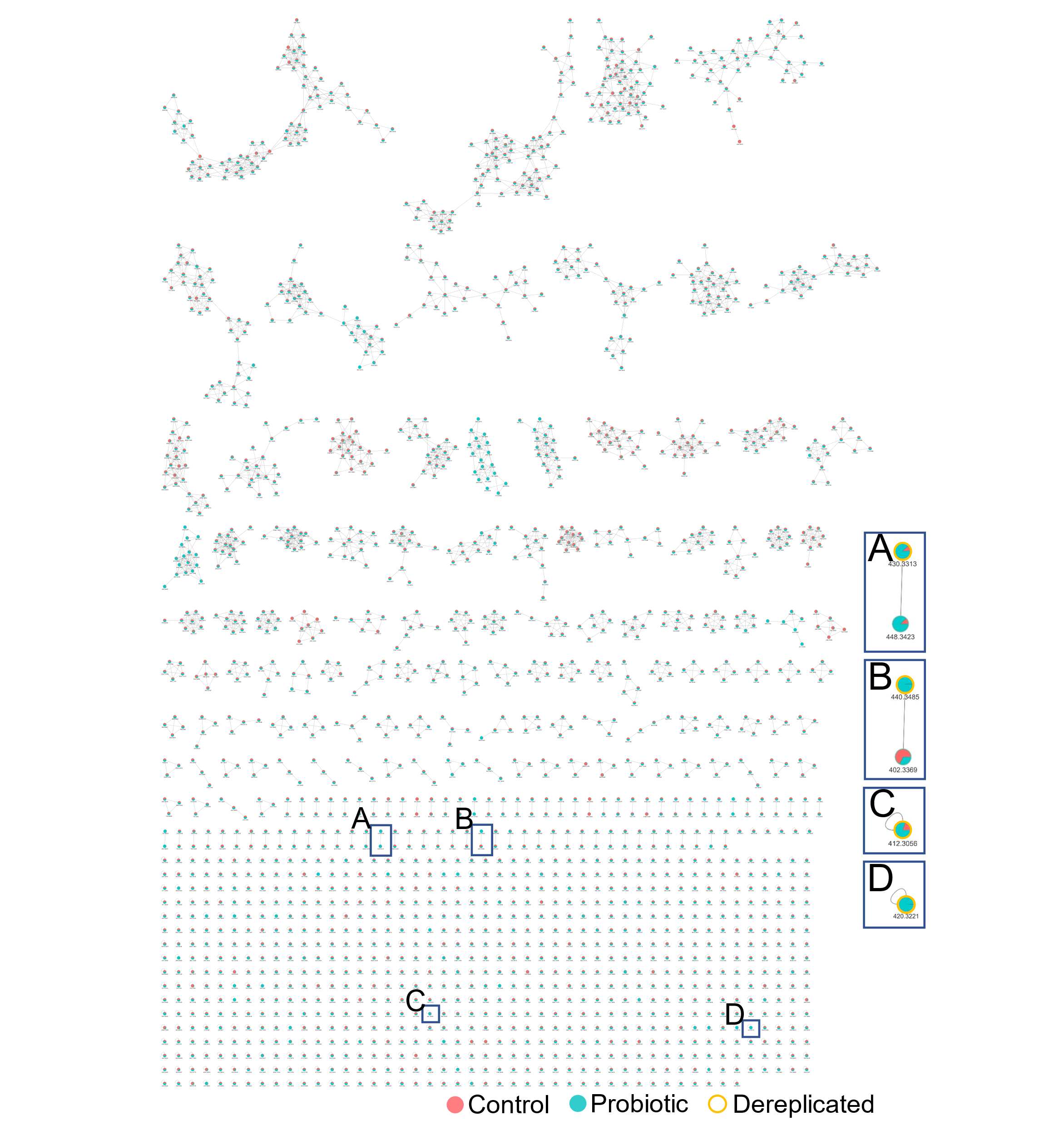

**Supplementary figure 6: Feature-based molecular network (FBMN) of the ions detected in the positive polarity (LC-MS/MS)**

Every node of the network represents a mass feature with a specific retention time, and is labelled with the *m/z* of the parent ion. The mass features are clustered together based on similar fragmentation patterns in their MS/MS spectrum. A pie chart is mapped to the nodes that shows the abundance of each feature in the control (salmon) and probiotic-supplemented (blue) samples. Dereplicated features are highlighted in yellow in the enlargements (A-D). **A** Phe-C18:1 (*m/z* 430.3313) **B** Citrulline-C18:1 (*m/z* 440.3485) **C** Glu-C18:1 (*m/z* 412.3056) **D** His-C18:1 (*m/z* 420.3321). The FBMN was illustrated using Cytoscape.

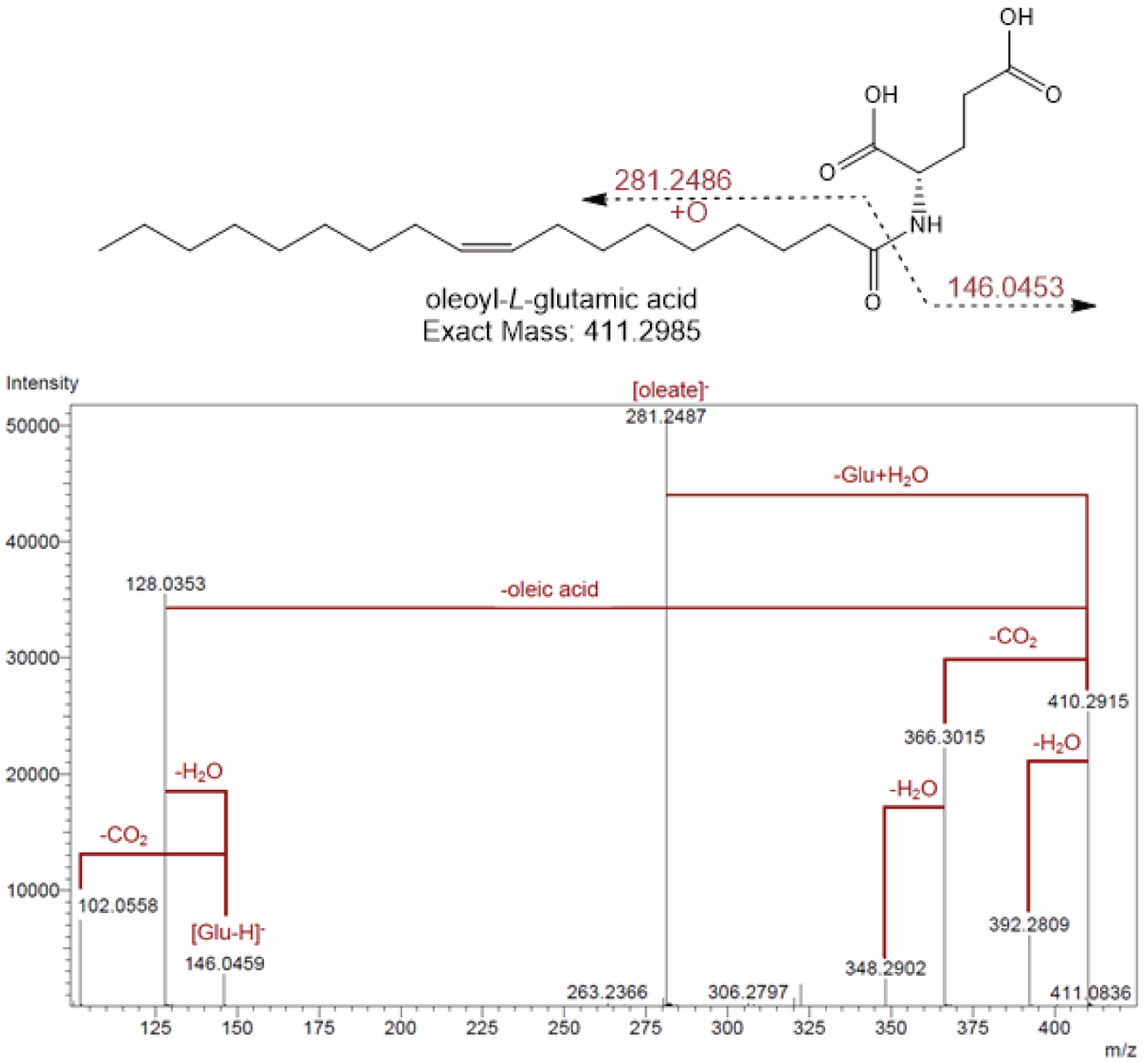

**Supplementary figure 7: Proposed molecular structure and MS/MS spectrum of the precursor mass *m/z* 410.2914 (RT 10.62 min) in the negative polarity**

Characteristic fragments and their calculated ion masses are indicated in the molecular structure. Neutral losses are highlighted in the MS/MS spectrum. Based on the fragmentation pattern, the mass feature was dereplicated as the fatty acid-amino acid conjugate Glu-C18:1.

**References**

1.	de Vos WM, Tilg H, van Hul M, Cani PD. Gut microbiome and health: mechanistic insights. *Gut* 2022; **71**: 1020–1032.

2.	Yu Z, Morrison M. Improved extraction of PCR-quality community DNA from digesta and fecal samples. *Biotechniques* 2004; **36**: 808–812.

3.	Lax S, Smith DP, Hampton-Marcell J, Owens SM, Handley KM, Scott NM, et al. Longitudinal analysis of microbial interaction between humans and the indoor environment. *Science (1979)* 2014; **345**: 1048–1052.

4.	Hasan NA, Young BA, Minard-Smith AT, Saeed K, Li H, Heizer EM, et al. Microbial community profiling of human saliva using shotgun metagenomic sequencing. *PLoS One* 2014; **9**.

5.	Ponnusamy D, Kozlova E v, Sha J, Erova TE, Azar SR, Fitts EC, et al. Cross-talk among flesh-eating Aeromonas hydrophila strains in mixed infection leading to necrotizing fasciitis.

6.	Ottesen A, Ramachandran P, Reed E, White JR, Hasan N, Subramanian P, et al. Enrichment dynamics of Listeria monocytogenes and the associated microbiome from naturally contaminated ice cream linked to a listeriosis outbreak. *BMC Microbiol* 2016; **16**.

7.	Nurk S, Meleshko D, Korobeynikov A, Pevzner PA. MetaSPAdes: A new versatile metagenomic assembler. *Genome Res* 2017; **27**: 824–834.

8.	Cantalapiedra CP, Hern̗andez-Plaza A, Letunic I, Bork P, Huerta-Cepas J. eggNOG-mapper v2: Functional Annotation, Orthology Assignments, and Domain Prediction at the Metagenomic Scale. *Mol Biol Evol* 2021; **38**: 5825–5829.

9.	Huerta-Cepas J, Szklarczyk D, Heller D, Hernández-Plaza A, Forslund SK, Cook H, et al. eggNOG 5.0: a hierarchical, functionally and phylogenetically annotated orthology resource based on 5090 organisms and 2502 viruses. *Nucleic Acids Res* 2019; **47**: D309–D314.

10.	Quinlan AR, Hall IM. BEDTools: a flexible suite of utilities for comparing genomic features. *Bioinformatics* 2010; **26**: 841–842.

11.	Ritchie ME, Phipson B, Wu D, Hu Y, Law CW, Shi W, et al. limma powers differential expression analyses for RNA-sequencing and microarray studies. *Nucleic Acids Res* 2015; **43**: e47.

12.	von Meijenfeldt FAB, Arkhipova K, Cambuy DD, Coutinho FH, Dutilh BE. Robust taxonomic classification of uncharted microbial sequences and bins with CAT and BAT. *Genome Biol* 2019; **20**: 1–14.

13.	Pluskal T, Castillo S, Villar-Briones A, Orešič M. MZmine 2: Modular framework for processing, visualizing, and analyzing mass spectrometry-based molecular profile data. *BMC Bioinformatics* 2010; **11**: 1–11.

14.	Xia J, Psychogios N, Young N, Wishart DS. MetaboAnalyst: a web server for metabolomic data analysis and interpretation. *Nucleic Acids Res* 2009; **37**: W652.

15.	Wang M, Carver JJ, Phelan V v., Sanchez LM, Garg N, Peng Y, et al. Sharing and community curation of mass spectrometry data with Global Natural Products Social Molecular Networking. *Nature Biotechnology 2016 34:8* 2016; **34**: 828–837.

16.	Nothias LF, Petras D, Schmid R, Dührkop K, Rainer J, Sarvepalli A, et al. Feature-based molecular networking in the GNPS analysis environment. *Nature Methods 2020 17:9* 2020; **17**: 905–908.
